# Adaptation to drought is coupled with slow growth in marginal silver fir (*Abies alba* Mill.) populations

**DOI:** 10.1101/531806

**Authors:** Katalin Csilléry, Nina Buchmann, Bruno Fady

**Author notes:** **Corresponding author:** Katalin Csilléry, Department of Evolutionary Biology and Environmental Studies, University of Zürich, Winterthurerstrasse 190, 8057 Zürich, Switzerland &.

## Abstract

Drought is increasingly considered as the most important selection pressure for forest trees in the context of climate change. We studied adaptation to drought in marginal populations of silver fir (*Abies alba* Mill.) from the French Mediterranean Alps. Drought tolerance was assessed using proxies both from seedlings and adult trees. We measured water stress response, growth and bud break of seedlings originating from 16 populations in a greenhouse common garden experiment (N=8199) and water use efficiency via *δ*^13^*C* of adult trees of the source populations *in-situ* (N=315). Further, 357 single nucleotide polymorphisms (SNPs) were used to uncover the demographic history of the populations. Demographic distances between populations were used to generate a null expectation for trait divergence, thereby detect the signature of natural selection. We found evidence for adaptive population divergence in drought tolerance across life stages. Seedlings originating from source populations with low soil water capacity resisted better to water stress in the greenhouse, and additionally, adult trees from these populations had a higher water use efficiency. Seedling growth showed an evolutionary trade-off with drought tolerance: seedlings with fast growth and high stature came from populations that had lower drought tolerance. In contrast, population divergence in bud break showed only a weak signal of adaptation, which was independent of that in drought tolerance. Variation in phenology between populations was associated with variance in temperature and drought frequency and severity at the source populations. Our results highlight the adaptive value of marginal populations, advance our understanding of the different processes that have allowed silver fir to cope with drought stress under a warming climate, and contribute to our knowledge to advise assisted migration programs.

## Introduction

Drought is one of the most important selection pressure for terrestrial plants, including forest trees. Recent evidence suggests that forest tree populations worldwide respond to the combination of global warming and drought, so-called global change-type drought, with increased mortality (Asner *et al*., 2016; Allen *et al*., 2015; McDowell *et al*., 2011). Hotter droughts also foster the appearance of other detrimental agents such as pests (pathogens, herbivores) or physical disturbances such as storms, leading to positive feedback loops that can further contribute to forest damage (*e.g*. Csilléry *et al*., 2017; Seidl *et al*., 2017). Die-back events of adult trees lead to a dramatic loss of biomass, which are increasingly raising alarms due to their effects on ecosystem functioning, carbon balance, and economy (Trumbore *et al*., 2015; Anderegg *et al*., 2015). However, forest decline can also go undetected when focusing on adult trees only. Indeed, the lack of sufficient seedling recruitment, the so-called extinction debt (Hanski & Ovaskainen, 2002), is likely a major issue in many forest tree species facing climate change (Talluto *et al*., 2017). Thus, in order to understand and predict the effect of climate change on forest ecosystems, and to plan adaptive forest management strategies, drought sensitivity has to be assessed across different developmental stages, from seedling to adults (McDowell *et al*., 2013).

Trees are sessile and long-lived organisms, and as such, they must cope with local water stress conditions, and its temporal fluctuations. As a result trees acquired a large variety of mechanisms to reduce the negative impacts of drought encompassing changes from micro, cellular or organ level changes, such as stomatal closure, to macro scales, such as phenological shifts (Moran *et al*., 2017). Indeed, water stress response can be best considered a collection of functional traits (a syndrome) that are more or less strongly correlated with each other due to genetic coupling (pleiotropy) and developmental constraints (Juenger, 2013). The physiology of adult tree death likely involves failures of the coupled hydraulic system and carbon dynamics, while the relative importance of the two often remains unclear (McDowell *et al*., 2011; Sala *et al*., 2012). For example, in southern, marginal silver fir and Scots pine populations, early wood tracheids in declining trees showed smaller lumen area with thicker cell wall, inducing a lower theoretical hydraulic conductivity (Pellizzari *et al*., 2016). While, in piñon pine, it was shown that both hydraulic failure and carbon starvation can lead to tree death (Sevanto *et al*., 2014).

Although selection occurs throughout the whole life cycle, early life stages play a critical role. The selective survival of drought resistant seedlings could be an efficient strategy if their is a genetic correlation between relevant seedling and adult drought tolerance traits (Donohue, 2014). For growth, genetic correlation across life stages has been documented. For example, Surles *et al*. (1993) found that embryo and seed weight predicted field breeding values for 5- and 15-year volume growth in slash pine (*Pinus elliottii*). These results highlight that studying seedlings can, in some cases, advance our understanding of adaptation in natural populations of long-lived organisms. For pleiotropy of drought tolerance traits there is less information on trees, but for example in *Arabidopsis*, a positive genetic correlation was found between water use efficiency calculated as *δ*^13^*C* and phenology, which was considered as drought-escape strategy (Mckay *et al*., 2003).

Marginal populations, especially those at the rear edge of the species distribution area, are often living descendants of glacial refugium populations (Hampe & Jump, 2011). It has long been recognized that such climate relicts can have high levels of neutral genetic diversity. However, it has often been overlooked that may also have accumulated adaptive genetic hence phenotypic variation during their evolutionary trajectories dominated by climate warming since the last glacial maximum (Sexton *et al*., 2009; Hampe & Jump, 2011). Indeed, several common garden studies of forest tree species report that seedlings from southern and drier provenances have higher drought tolerance than northern more humid populations, for example, in European beech (Robson *et al*., 2012; Kreyling *et al*., 2014) or in cork oak (*e.g*. Ramírez-Valiente *et al*., 2010, 2018). Southern populations have probably been exposed to stronger and longer-lasting climate-related selection than their conspecifics from the main distribution range, thus they can be extremely useful resources for assessing the adaptive capacity of populations to ongoing climate change (Fady *et al*., 2016).

Drought will be a primary threat to forest ecosystems all over Europe during the 21^*st*^ century, but in particular, in the Mediterranean, where climate models are concordant to predict a pronounced decrease in precipitation and an increase in temperature, principally during the growing season (Giorgi & Lionello, 2008; Cramer *et al*., 2018). Here, we study marginal populations of an ecologically and economically important, late successional European conifer, silver fir (*Abies alba* Mill.) from the French Mediterranean Alps. In this region the species distribution is predominantly limited by water availability and populations are recurrently exposed to drought conditions (Roschanski *et al*., 2016; Cailleret & Davi, 2011). Further, the study area is characterized by a high degree of environmental heterogeneity, where climatic conditions may rapidly change from hot, thermo-Mediterranean to colder and moister mountain-Mediterranean (Sagnard *et al*., 2002). Thus, in this region, silver fir is often found in rather unusual habitats for the species, e.g. on north facing slopes of isolated mountains surrounded by vineyards. These suitable habitats mostly found at elevations between approximately 900 and 1600 m above sea level.

The demographic history of silver fir in the French Mediterranean Alps has been largely debated. Palaeobotanical and DNA marker data suggest that silver fir in Europe recolonized its current habitats after the last glacial maximum from five refugia, and populations from the French Mediterranean Alps most likely originate from a refugium in Central Italy (Liepelt *et al*., 2009). More recently, Ruosch *et al*. (2016) used a dynamic global vegetation model validated using historical pollen data, and found that silver fir could have maintained populations in the French Mediterranean Alps during the last glacial maximum about 21 kya ago. Then, approximately 15 kya ago populations in southern France went through a strong bottleneck, but recovered from it and since then populations are slowly declining. Although the study of Ruosch *et al*. (2016) supports the idea that the southern French populations survived several climatic changes and likely have accumulated adaptations, their origin remains unclear as it dates back to earlier times than the last glacial maximum.

In this study, we investigate the evolution of drought tolerance across 16 marginal populations from the French Mediterranean Alps. We combine measures from seedlings grown in a greenhouse common garden experiment with a proxy of water use efficiency, *δ*^13^*C*, from adult trees *in-situ* to contrast the drought tolerance different populations. Growth and spring bud-break phenology recorded on the same seedlings allows us to explore the trait space for correlated character evolution and trade-offs. Using genetic marker data, we test the hypothesis if phenotypic differentiation between populations is the result of demography, i.e./ genetic drift, or natural selection. This distinction is particularly important in this region because we are studying small and isolated populations, where genetic drift could have also led to differences between population (Whitlock & Guillaume, 2009). Indeed the sampling region is dominated by the presence of small isolated population patches, including an island population, Corsica. Finally, we explore a large range of environmental variables, including topography, soil and climate to identify potential drivers of adaptive population divergence to drought.

## Material and Methods

### Study system and sampling

Silver fir is an ecologically and economically important European forest tree species that can tolerate episodes of drought thanks to its deep rooting system (*e.g*. Lebourgeois *et al*., 2013; Vitali *et al*., 2017) or indirectly, due to its high tolerance to bark beetle attacks (Wermelinger, 2004). We studied 16 silver fir stands across the French Mediterranean Alps (Fig. 1a, Table S1). Sampling sites were chosen within an elevational range representative of the local silver fir forests, isolated trees were avoided. In the fall of 1994, seeds were collected from 19–43 dominant trees from each site, respecting a minimal distance of 30 m between trees (Sagnard *et al*., 2002). The weight of 1000 seeds was measured for each mother tree and used to control for maternal effects. The sites were re-visited in 2016, when needle samples were collected from approximately 20 trees at each site. In a few cases, we were able to re-identify the mother trees, which we sampled. Otherwise, we sampled other mature trees we found respecting at least 100 m distance between them. One hundred meters are generally sufficient to sample unrelated trees (Mosca *et al*., 2018), which was necessary to obtain unbiased estimates of the population allele frequencies.

**Fig. 1:**
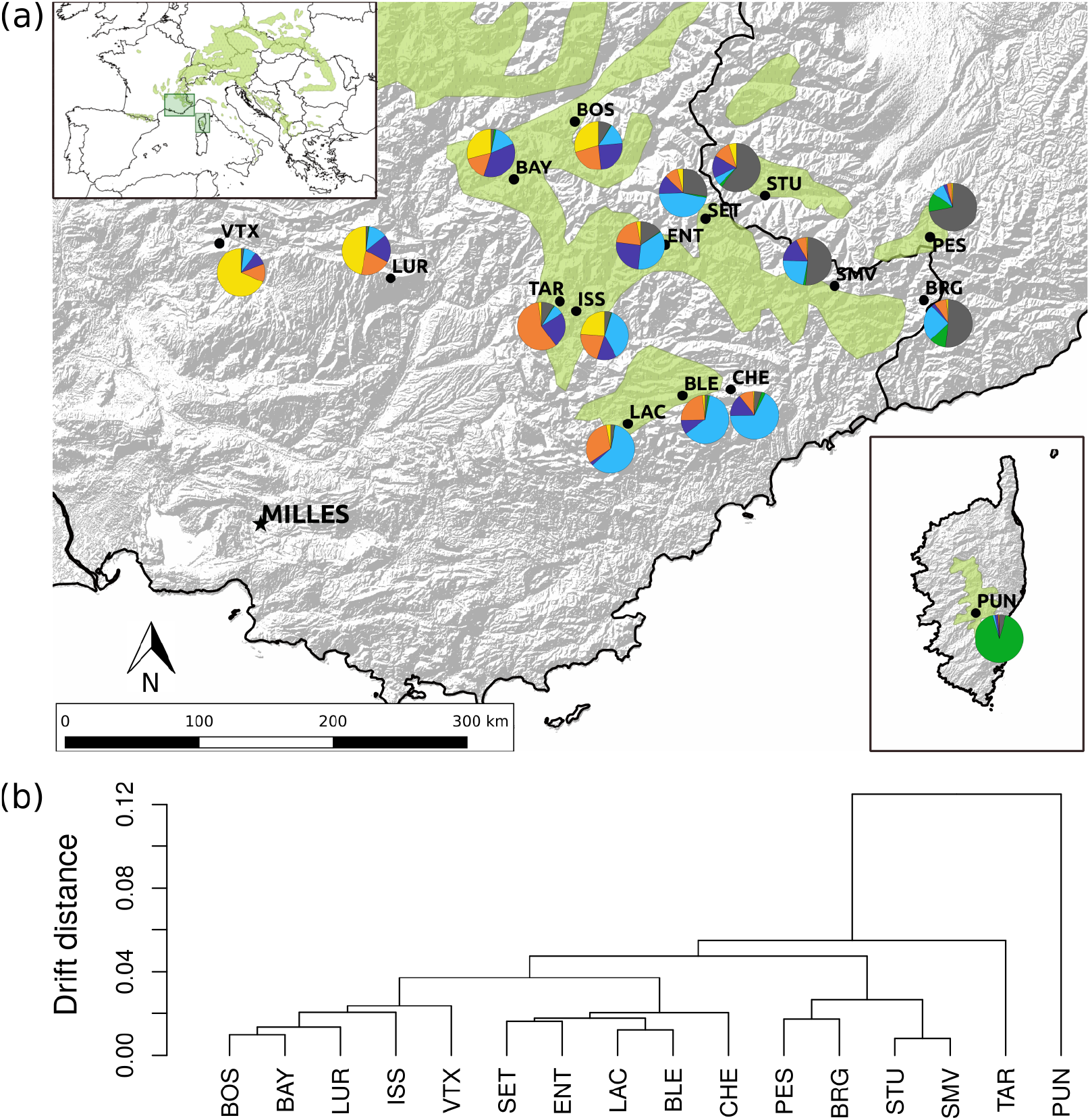
Population genetic structure of 16 silver fir (*Abies alba*) populations from the French Mediterranean Alps inferred from 357 SNP loci. **(b)** Geographic location of the sampling sites and proportion of individuals belonging to each of the six genetic clusters identified using the admixture model of the software *Structure*. **(b)** Neutral demographic, so-called drift distances between populations estimated using the admixture F-model implemented in the R package *RAFM*.

### Common garden experiment

In 1995, seeds were sown in an experimental forest nursery located in Milles, near Aix-en-Provence in southeastern France (Table 1). Seedlings were grown in 600 cm^3^ containers, grouped in plastic crates by sets of 32. Each mother tree (subsequently family) was represented by 16–20 seedlings. The experimental design consisted of two greenhouses each divided into two complete blocks. Populations, families and seedlings were randomized across greenhouses and blocks. Greenhouse 2 was exposed to a water stress treatment that involved a complete lack of watering starting from 6 May 1998. Greenhouse 1 received watering for the whole duration of the experiment.

**Table 1:**
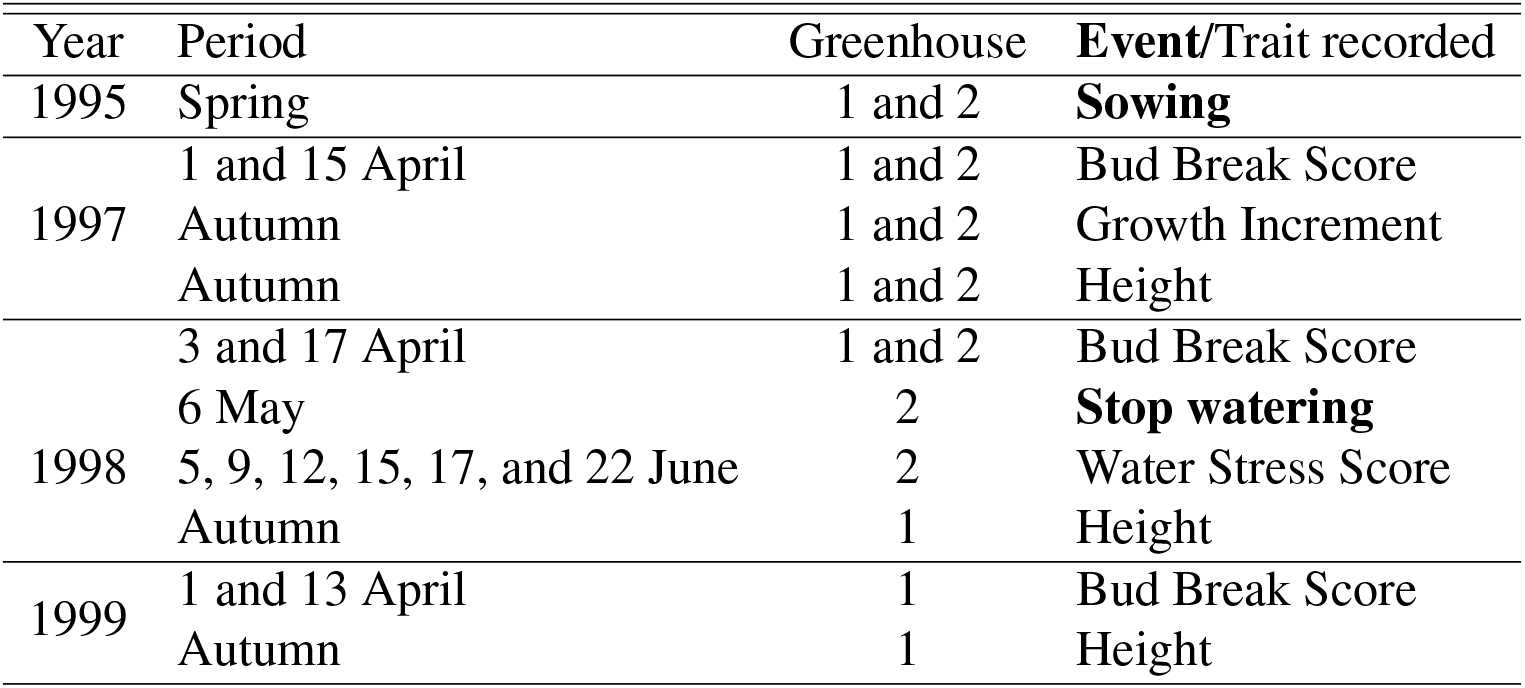
The time-line of the common garden experiment performed in the experimental forest nursery located in Milles (see Fig. 1). See Table 2 for definitions of the traits.

Traits were recorded starting from the 2^*nd*^ growing season (i.e. from 1997) to the 4^*th*^ (i.e. until 1999) (Table 1). We analyzed a total of 8199 observations, of which 3931 were in greenhouse 1 and 4267 in greenhouse 2. Growth Increment and Height were used as raw measurements (trait names are capitalized hereafter; see Table 2 for trait definitions). Spring bud break phenology was scored at two dates each year from which we calculated a Bud Break Score (see Table 2). Bud Break Score ranged between two and 10, where higher numbers indicate earlier bud break. Response to water stress was scored at five consecutive dates in 1998 (Table 1). We used a derived trait, Water Stress Score, to characterize response to water stress, which ranged between zero and 10 (see Table 2). Low values indicate drought-hardy seedlings and high values indicate drought sensitive seedlings. The distribution of the Water Stress Score was highly skewed and zero inflated (see Water Stress Score (raw sum) on Fig. S1). Thus, we calculated an integrative measure of water stress by weighting the stress scores with the log Julian dates of the observations. This weighted Water Stress Score had a close to Normal distribution (see Water Stress Score on Fig. S1), and accounted for the evolution of the response to stress by giving higher weight to earlier responses to stress (i.e. to the most sensitive seedlings).

**Table 2:**
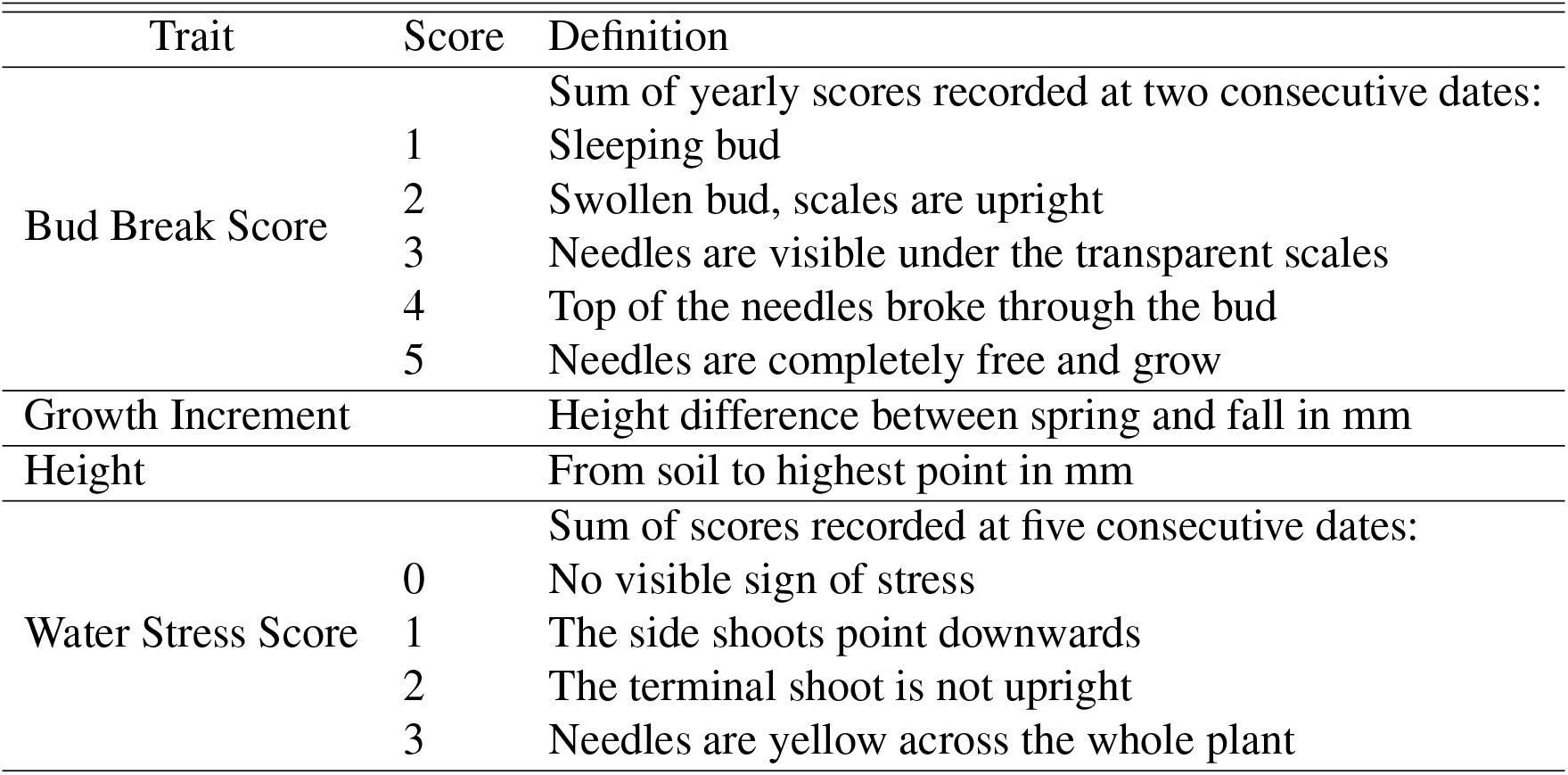
Silver fir (*Abies alba* Mill.) seedling traits measured in a common garden and their definitions.

### Adult tree data

Fresh needles from adult trees were lyophiliozed for at least 48 h within a few days of their collection and sent for DNA extraction and genotyping to LGC Genomics (Middlesex, United Kingdom). A total of 315 trees were genotyped at 357 single-nucleotide polymorphism (SNP) loci using KASP arrays and the all-inclusive service from LGC Genomics (Middlesex, UK). We used 267 SNPs from Roschanski *et al*. (2016) and additionally selected 110 new putatively neutral SNPs from the transcriptome assembly of Roschanski *et al*. (2016), based on respective values of Tajima’s D between 2 and -2 and dN/dS between 0.9 and 1.1, and with low LD with the existing SNPs (*r*^2^ <0.1 and p-value > 0.05). Further, we selected 149 SNPs from the control panel of Mosca *et al*. (2012) that had less than 5% missing data in that study. Three out of the four populations studied in Roschanski *et al*. (2016) were included in our set of 16 populations.

*δ*^13^*C* of 165 trees adult trees, i.e. ten trees per population on average, was measured at the ETH Grassland Sciences Isolab. Approximately 80 mg lyophilized needle material was milled in 2 ml polypropylene tubes equipped with a glass ball (diameter of 5 mm) (4min at 30 Hz). Milled samples were directly weighted in small tin capsules (approx. 5 mg, XPR2 microbalance from Mettler Toledo). Sub-samples of approximately 5 mg needle powder were combusted in an elemental analyzer (Flash EA by Thermo Finnigan, Bremen, Germany) coupled to an isotope ratio mass spectrometer (Delta XP by Thermo Finnigan, Bremen, Germany) by a Conflo II interface (Thermo Finnigan, Bremen, Germany).

### Environmental data

We used downscaled historical climatic data to characterize environmental conditions at each sampling site. In order to obtain the closest representation of the climate for the period when the current populations were established, we used data from 1 January 1901 to 31 December 1978. The choice of this period was justified by two facts: (i) no observation-based climate data go back further in time, and (ii) starting from approximately 1980, the temperature time series are overwhelmed by the effect of global warming (Harris *et al*., 2014). We used the delta method for statistical downscaling (Hay *et al*., 2000) to obtain mean monthly temperature and precipitation on a 1 km grid scale. The reference climatic data set for downscaling was the 0.5° resolution CRU TS v. 4.01 data (20 September 2017 release, Harris *et al*. (2014)), while the downscaling was based on the overlapping period with the 1 km resolution Chelsa data (Karger *et al*., 2017).

We calculated several bioclimatic indexes to characterize the environment at each site, including the 19 standard bioclimatic variables (Booth *et al*., 2014) implemented in the R package *dismo* (Hijmans *et al*., 2017), two potential evapotranspiration (PET) indexes and a standardized precipitation - evapotranspiration index (SPEI) using the R package *SPEĪ* (Beguería & Vicente-Serrano, 2017), and two indicators of late frost (Table S2). From the yearly time series of SPEI, we calculated indices of drought severity and frequency over the whole period of 78 years (Table S2). Available Water Capacity of the soil (volumetric fraction) until wilting point was obtained at a 250 m resolution from the SoilGrids250 data base for depth 5, 15, 30 and 60 cm (Hengl *et al*., 2017). 5 and 15 cm, and 30 and 60 cm were strongly correlated, so we averaged these two depths.

### Statistical analysis

#### Population structure and demography

First, we used the Bayesian clustering algorithm implemented in the software *Structure* v.2.3.4 (Pritchard *et al*., 2000) to find clusters of genetically related individuals across the sampled populations using the SNP data. We used the admixture model with correlated allele frequencies following Falush *et al*. (2003). Further, we included sampling location information to improve clustering performance (“locprior model”, Hubisz *et al*. (2009)). We estimated the prior population allele frequency parameter (λ) from the data to account for the fact that SNPs often have rare minor alleles. We estimated λ using K=1 to avoid non-identifiability with the other hyperparameters (λ, *α*, F). λ was consistently around 0.65 across ten repeated runs (range: 0.63-0.66, median: 0.65). Then, we tested K values from 1 to 19 using ten independent Markov chains for each K, and 500,000 burn-in iterations and 500,000 iterations for estimation of the membership coefficients. Different numbers of clusters (K) were compared with *StructureHarvester* (Earl *et al*., 2012) using the *LnPr*(*X|K*) and Evanno *et al.’s* (2005) method. Admixture coefficients were averaged across ten repeated runs using CLUMPP v.1.1.2 Jakobsson & Rosenberg (2007) using an exhaustive search for K≤3, the Greedy algorithm for any K>3, and large-K-Greedy for K≥5.

Second, we estimated the demographic distances between populations from variation in the SNP allele frequencies. For this, we used an admixture F-model (AFM), which assumes that populations have diverged from a common ancestral pool due to genetic drift (Karhunen & Ovaskainen, 2012). This model can disentangle between small population size and isolation, which are the principal mechanisms underlying drift. We estimated these so-called drift distances, a.k.a. the coancestry matrix, between all pairs of populations using Metropolis-Hastings algorithm implemented in the R package RAFM^1^ (Karhunen & Ovaskainen, 2012). We ran 10 independent Markov chains with a burn-in of 20,000 iterations followed by 10,000 iterations for estimation with a thinning interval of 10, and averaged the posterior estimates of the coancestry matrix across the chains.

#### Variance components and evolutionary potential

We used a mixed-effects model defined at the level of each individual (seedling), called the animal model (*e.g*. Wilson *et al*., 2010), to partition the trait variance to different components. The advantage of the animal model is that it gives a direct estimate of the trait variance due to genetic factors, the additive genetic variance (*V_A_*). We ran a separate model for each trait, where block was included as a fixed effect (with two or four values depending if one or both greenhouses were considered), and seed weight, population, and seedling as random effects, and a random residual error for each seedling. *V_A_* was estimated from the covariance between relatives, for which, we reconstructed a pedigree assuming seedlings from the same mother tree to be half-sibs. We implemented the model using the R package *ASreml-R* 3.0, and constructed the inverse kinship matrix using the function *asreml.Ainverse*.

We performed several tests to check the validity of the above full model. We tested if environmental heterogeneity generated by the experimental designed contributed to the trait variance by comparing models with and without block effect using a Wald-test (*wald.asreml*). We also tested for the significance of the random effects using a likelihood ratio test between models with and without seed weight and population, and also tested significance of including the pedigree itself (referred to “family” effect). Finally, the above model assumes that the 16 populations have a common *V_A_*. This assumption may not hold because the populations were selected from an environmentally and perhaps demographically heterogeneous region. In order to test the hypothesis of heterogeneity in *V_A_* across the study region, we repeated the above analysis for the three main genetic clusters that the above demographic analysis identified. The three clusters were: BOS, BAY, LUR, ISS, VTX (North-Western cluster, 2496 observations); SET, ENT, LAC, BLE, CHE (Central cluster, 2639 observations); and PES, BRG, STU and SMV (Eastern clusters, 1883 observations). TAR and PUN were excluded from this regional analysis because they did not belong to any of the three clusters (Fig. 1).

Since the variance components are largely unaffected by the inclusion of model terms that do not contribute to the trait variance, we used the full model to estimate the *V_A_* for each trait. From these, we derived two commonly used measures of the evolutionary potential: the *V_A_* standardized by the total variance, i.e. the heritabilty (*h*^2^, Falconer & Mackay (1996)) and by the mean trait value, which is the additive genetic coefficient of variation (*CV_A_*, Houle (1992).

#### Adaptive divergence and environmental drivers

We used the test of adaptive divergence (*S*-test) developed by Ovaskainen *et al*. (2011) implemented in the R package *driftsel*^2^ to distinguish between the effects of drift and natural selection in shaping the trait values across the 16 populations. *driftsel* also implements an animal model. However, it not only accounts for the pedigree, but also the demographic distances between the populations estimated from genetic marker data (see above). To achieve this, *driftsel* coestimates the additive genetic mean and variance of the trait value in an assumed ancestral population and the deviations from the ancestral mean for each population, i.e. the population effects. Covariates can also be incorporated, and here, we used block and seed weight. The *S*-statistic is used to summarize the evidence for adaptive divergence across all populations. *S* = 0.5 indicates consistency with neutrality, S = 0 implies a match with purifying, and *S* =1 with diversifying selection. The S-statistic cannot distinguish between many, slightly diverged, populations or a single or few greatly diverged populations. Thus, following Csilléry *et al*. (2018), we also summarized the evidence for adaptation for each population. We will conclude that a particular population diverged more from the ancestral mean than expected by drift if the 95% of the posterior distribution of the mean additive trait value is outside of the drift envelop. Finally, note that *driftsel* uses a Bayesian approach, so we will use “unusual” and not “significant” to indicate results with strong evidence of adaptive divergence.

We ran three independent chains for each trait with a burn-in of 100,000 iterations followed by 30,000 iterations for parameter estimation and using a thinning interval of 10. We ran separate analysis for the two greenhouses mainly because the Markov chain Monte Carlo analysis of *driftsel* was not computationally feasible for the full data set of nearly 8000 observations. Although *driftsel* can accommodate multi-trait tests of selection, we were unable to reach convergence for trait-pairs given the size of the data set (tried up to 200,000 iterations, which involves a run time of over 4 weeks; results not shown). In order to summarize adaptation in a multi-trait space, we performed a Principal Component Analysis of the deviations of mean additive trait values from the estimated ancestral mean standardized by the trait variance expected based on drift.

In order to identify environmental variables that might have imposed selection pressure on particular populations, we used the *H*\*-test introduced by Csilléry *et al*. (2018), which is a standardized version of the *H*-test Karhunen *et al*. (2014). The *H*\*-test is used to evaluate if environment is correlated with the estimated trait divergence, and the *H*\* statistic has a similar interpretation to the *S* statistic. To avoid multiple testing of the large number of environmental variables (Table S2), we reduced the environmental variables to synthetic variables using a Principal Component Analysis (PCA) using the *prcomp* function in R. All variables were centered and scaled. The first four PC axes explained 88.9% of the variance. Based on the variables with the highest loadings on each of the PC axes, the four PCA variables will be referred to as Temperature variance (PC1), Mean temperature (PC2), Soil water capacity (PC3), Drought (PC4); see Table 3.

**Table 3:** Principal Component Analysis (PCA) of 33 environmental variables, see Materials and Methods for details the analysis and a full list of variables in Table S2. The first four PC axes explained 88.9% of the variance in the environmental variables. The first three environmental variables with the highest loadings on PC1 to PC4. Column names indicate synthetic names and variance explained for the PC axes. Abbreviations: T: Temperature, P: Precipitation AWC: Available Water Capacity, PET: Potential Evapotranspitation; SPEI: Standardised Precipitation-Evapotranspiration Index.

| Temperature variance<br>PC1 (40.1%) |  | Mean temperature<br>PC2 (22.2%) |  | Soil water capacity<br>PC3 (17.8%) |  | Drought<br>PC4 (8.8%) |  |
| --- | --- | --- | --- | --- | --- | --- | --- |
| Variables | Loadings | Variables | Loadings | Variables | Loadings | Variables | Loadings |
| T Annual Range | -0.28 | Late frost | 0.31 | AWC 30-60cm | 0.36 | N months with SPEI < -2 | 0.49 |
| Mean Diurnal Range | -0.28 | PET | 0.3 | AWC 5-15cm | 0.29 | 5% quantile of SPEI | -0.35 |
| T Seasonality | -0.28 | Mean Annual T | 0.29 | Mean Annual T | 0.27 | P of coldest Quarter | 0.33 |

## Results

### Population structure and demographic history

Using *Structure*, we obtained the highest support for the presence of six (log-likelihood method) and four (Evanno method) genetic clusters across the 16 silver fir populations (Fig. S2). The population from the island of Corsica (PUN) was the most different from the other populations; in fact, a separation between Corsican and mainland populations was already clear with K=2 (Fig. S2a). The genetic structure of the mainland populations was characterized by isolation-by-distance both from east to west and from south to north (Fig. 1a). Interestingly, the two populations that were the geographically closest (TAR and ISS) and situated in the middle of the sampling range, were one of the most strongly differentiated population pairs.

Convergence was good using *RAFM*, the mean potential scale reduction factor (Gelman & Rubin, 1992) was one across the ten chains and ranged between 0.9 and 1.1. Using the admixture F-model, results were similar to those obtained with *Structure* regardless the different model assumptions: *Structure* clusters individuals to an arbitrary number of groups, while *RAFM* assume the presence of single ancestral gene pool from which all populations had been derived while maintaing varying levels of gene flow. Fig. 1b shows that with *RAFM*, again PUN was the most distant population, followed by the separation of TAR. The other populations were grouped into three clusters that corresponded to geographic regions of nearby populations possibly connected by gene flow.

### Sources of trait variation and evolutionary potential

Comparison of animal models including different effects revealed that heterogeneity within the experiment captured by block had a significant effect on the variation of most traits (Table 4). For three traits, Bud Break Score 1997 and 1998, and Growth Increment 1997, we were able to compare the effect of the greenhouse. We found that variance components were similar when taking all data or just one of the greenhouses, with slightly higher estimates for one greenhouse only (Table 4, Fig. 2). The effect of year on the traits was addressed indirectly using Bud Break Score and Height that were measured across three years of the experiment in greenhouse 1. We found that seedlings most likely acclimated to the experimental conditions: the block effect disappeared for Bud Break Score 1998 and 1999 and for Height 1998. Further, we observed that the heritability decreased with years, most likely also due to acclimation, which made families more similar to each other (Fig. 2). However, it is also possible that maternal effects, other than seed weight, have contributed to an inflated heritability in the first year of the experiment, i.e. in 1997.

**Fig. 2:**
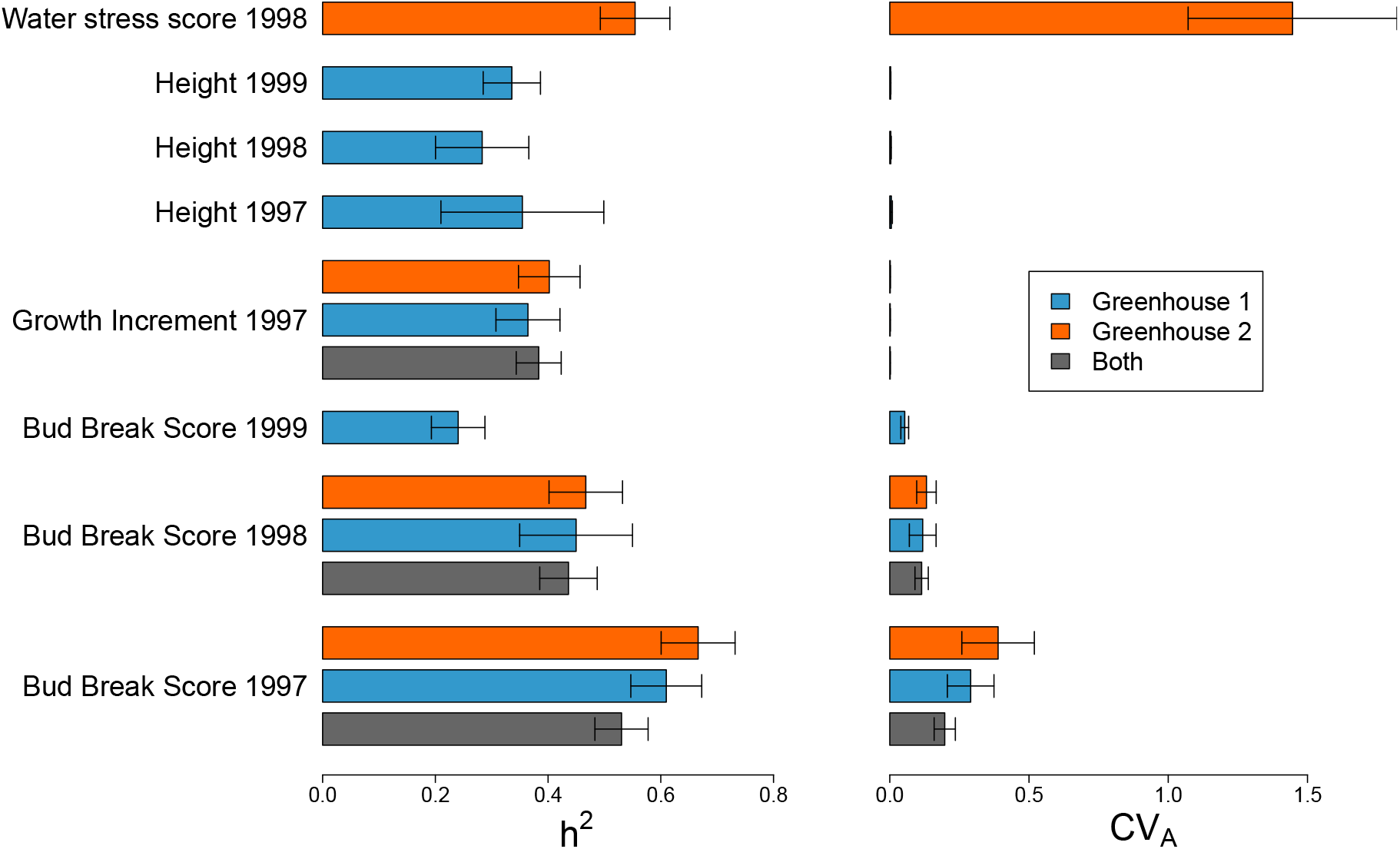
Heritability (*h*^2^) and additive genetic coefficient of variation (*CV_A_*) of 11 traits measured on silver fir (*Abies alba*) seedling in a common garden. The experiment consisted of two greenhouses and counted 8199 observations. Not all traits were scored in both greenhouses. Greenhouse 2 received a water stress treatment in 1998 after which only Water Stress Score was recorded. Parameters were estimated using an animal model implemented in *ASreml-R* including block as a fixed effect, and seed weight and population as random effects.

**Table 4:** Model comparison of 11 silver fir (*Abies alba*) seedling traits measured in a common garden using an animal model implemented in *ASreml-R*. Seedlings were grown in two greenhouses each comprising two blocks. Populations and families were randomized across greenhouses and blocks. N indicates the number the observations. The role of Block (fixed effect) was tested using a Wald-test, while the role of Family (i.e. the pedigree), and Seed weight and Population (random effects) were tested using a likelihood ratio test by excluding each of these variables one-by-one.

| Trait | Greenhouse | N | Block | Family | Seed weight | Population |
| --- | --- | --- | --- | --- | --- | --- |
| Bud Break Score 1997 | 1 | 3862 | 0.0020 | < 0.0001 | > 0.9999 | < 0.0001 |
|  | 2 | 3883 | < 0.0001 | < 0.0001 | > 0.9999 | < 0.0001 |
| Bud Break Score 1998 | 1 | 1566 | 0.2029 | < 0.0001 | > 0.9999 | 0.1356 |
|  | 2 | 2935 | 0.0017 | < 0.0001 | 0.3801 | < 0.0001 |
| Bud Break Score 1999 | 1 | 3592 | 0.3522 | < 0.0001 | 0.0784 | 0.0002 |
| Growth Increment 1997 | 1 | 3208 | < 0.0001 | < 0.0001 | < 0.0001 | < 0.0001 |
|  | 2 | 3620 | < 0.0001 | < 0.0001 | < 0.0001 | < 0.0001 |
| Height 1997 | 1 | 898 | 0.0006 | 0.0071 | 0.3228 | 0.0002 |
| Height 1998 | 1 | 1683 | 0.2588 | 0.0001 | 0.0482 | < 0.0001 |
| Height 1999 | 1 | 3705 | < 0.0001 | < 0.0001 | 0.0004 | < 0.0001 |
| Water Stress Score 1998 | 2 | 3620 | < 0.0001 | < 0.0001 | < 0.0001 | < 0.0001 |

Family (i.e. the pedigree) always explained a large percentage of the trait variation and a model with pedigree was always better than a model without it (Table 4). All traits expressed a high heritability, with the largest value observed for Bud Break Score 1997, especially in the Water Stressed blocks (0.69), while the lowest value, 0.24, was observed for Bud Break Score 1999. The additive genetic coefficient of variation (CVA) was the highest for the Water Stress Score, followed by Bud Break Score 1997, and it was negligible for Growth Increment and Height (Fig. 2). We tested for the effect of seed weight to assess if genetic or non-genetic maternal effects explain trait variation. Seed weight explained a negligible (at the maximum 0.027%, Fig. S3), yet significant part of trait variation in growth traits, i.e. Growth Increment and Height, but not in phenology (Table 4). Population of origin also explained a significant part of trait variation (Table 4). The proportion of variance explained by population varied between 9.8% (Height 1998) and 0.7% (Bud Break Score 1998) with a mean of 5% across traits (Fig. S3). Finally, the regional analysis, where we considered populations only from one of the demographically most uniform clusters, revealed the same relative differences between the traits in *h*^2^ and *CV_A_*, however, estimates were often higher (i.e. inflated) most likely owing to a smaller sample size (Fig. S4).

### Adaptive trait divergence and its environmental drivers

Trait divergence between populations that cannot be explained by neutral demographic processes (i.e. drift) alone is likely the result of adaptation. We detected adaptive divergence using the *S*-test in several traits (Fig. 3). Convergence was achieved using *driftsel* for all traits, the mean potential scale reduction factor (Gelman & Rubin, 1992) was 1.05 across the three chains and ranged between 0.85 and 1.15 across all traits. Overall, Water Stress Score and growth traits, i.e. Height and Growth Increment, showed stronger evidence of adaptive divergence (i.e. higher S values) than Bud Break Score (Fig. 3). In fact, spring phenology showed adaptive divergence in 1997, but the signal disappeared with time (S nearly 50% means no difference from neutrality). The environmental drivers of adaptation were also different among traits and showed a clustering into two distinct groups. While population divergence in phenology was mainly influenced by temperature variance (PC1), and drought (PC4), divergence in growth traits and Water Stress Score were influenced by mean temperature (PC2) and soil water capacity (PC3) (*H*-test, Fig. 3, Table 3).

**Fig. 3:**
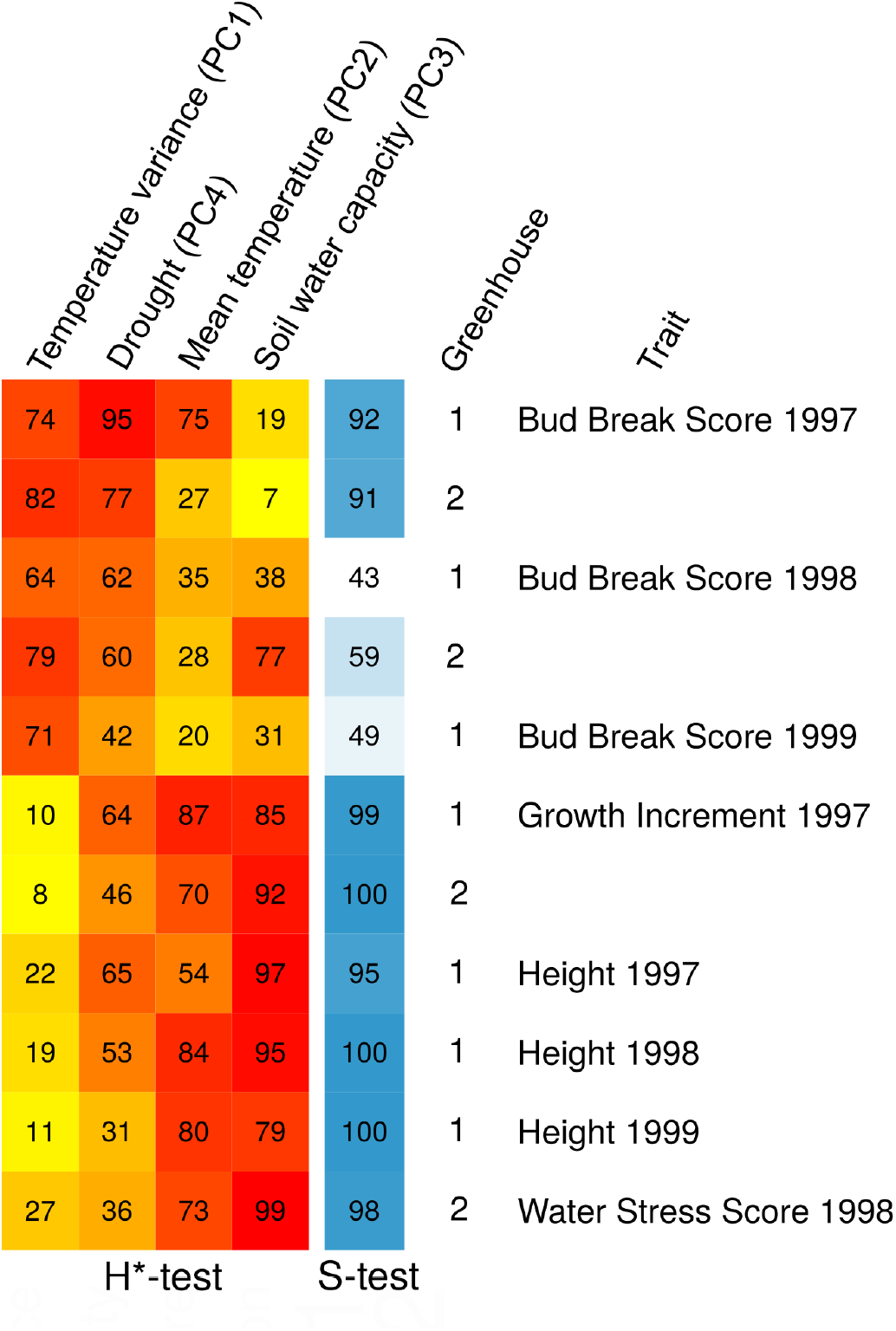
Evidence of adaptive divergence across 16 silver fir (*Abies alba*) populations at 11 traits measured on seedlings in a common garden. The *S*-test of adaptive divergence (Ovaskainen *et al*., 2011) and *H*\*-test of environmental drivers (Karhunen *et al*., 2014; Csilléry *et al*., 2018) are expressed as percentages. *S* and *H*\* closer to 1 (i.e. 100%) indicate higher evidence for adaptation, while values close to 0.5 (i.e. 50%) indicate neutrality. *H*\*-tests were performed on synthetic environmental variables derived using a Principal Component Analysis of variables listed in Table 3.

Next, we asked the question that in which directions populations evolved from a hypothetical ancestral trait value. For this, we used inferred the ancestral trait values and the expected level of trait divergence based on drift alone, and contrasted with the actual additive genetic trait values of the populations (Fig. 4a-d). We detected unusually early bud break for TAR, BAY, BOS and VTX in 1997 (Fig. 4a). Typical of a continental-type climate, these populations experienced the highest variance in temperature (PC1) at their environment of origin, the longest drought periods and winter precipitation in the form of snow (Table 3, Fig. 3). PUN, BRG, PES had the latest bud break, but their additive genetic trait values were within a range that could be expected based on neutral demographic processes alone (Fig. 4a). Nevertheless, their climate was the opposite of the populations with the earliest bud break, and characterized by low temperature variance and lack of prolonged drought periods (Table 3).

**Fig. 4:**
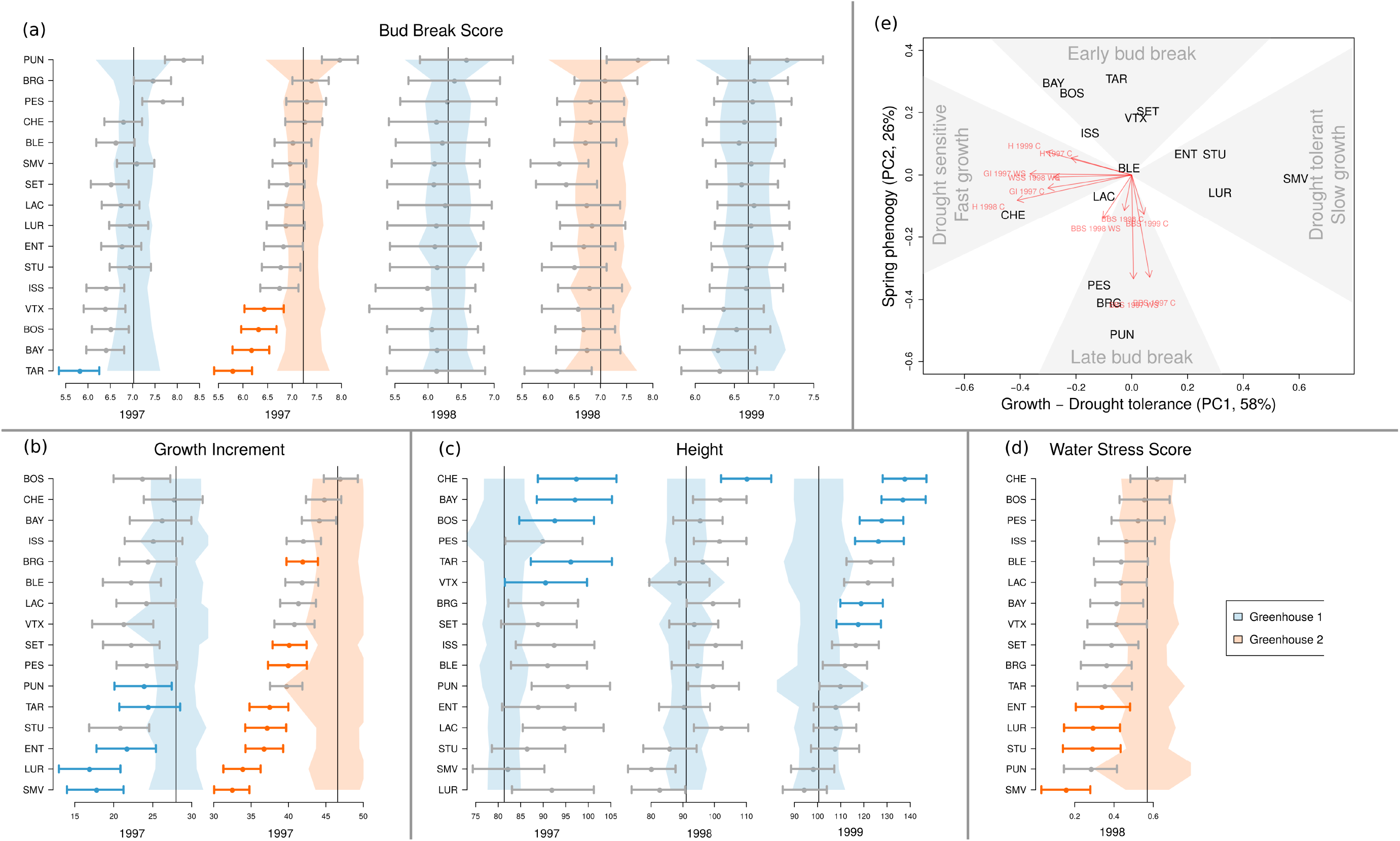
Evidence of adaptive divergence across 16 silver fir (*Abies alba*) populations at 11 traits measured on seedlings in a common garden. Each trait is shown in a panel **(a-d)**, grouping together its measures from different years and/or greenhouses. Each individual plot **a-d** shows the ancestral mean additive trait value (vertical line), the amount of trait divergence from this mean that is expected based on drift (light blue or orange envelop for greenhouse 1 and 2, respectively), and the mean and 95% credible interval of the posterior distribution of the additive trait values for each population (horizontal error bars) estimated using *driftsel*. The horizontal error bars are in blue or orange for populations with unusual divergence, which is indicative of adaptation; see Materials and Methods for details. Populations are ordered for each panel **a-d** according to the trait values for the year/greenhouse with the highest number of unusually diverged populations. **(e)** Summary of divergence across all traits using a Principal Component Analysis (PCA); see Materials and Methods for details. The first two PC axes are given synthetic names according to the variance they explained in the standardized trait divergence across the 11 traits.

Five to eight populations, depending on the greenhouse, showed unusually low Growth Increment: SMV, LUR, ENT, STU, TAR, PUN, PES, and SET (Fig. 4b), and one to six, depending on the year, populations unusually high Height: CHE, BAY, BOS, PES, TAR, VTX, BRG, and SET (Fig.4c). The fact that only one population showed unusual trait divergence for Height in 1998 could be due to the lower sample size in this year (see Table 4). we performed a Principal Component Analysis (PCA) of the standardized trait divergences to identify adaptive strategies in the whole trait space. The PCA analysis revealed that, not surprisingly, growth and height were strongly correlated (Fig 4e): populations that exhibited relatively fast growth also showed an unusually tall stature. The main environmental drivers of divergence in growth were soil water capacity (PC3) and mean temperature (PC2). Higher soil water capacity and warmer climate led to enhanced growth (Fig. 3, Table 3).

Four populations had unusually high levels of drought tolerance, i.e. lower Water Stress Score: SMV, STU, LUR, ENT (Fig. 4d). PUN had the second lowest Water Stress Score, but due to its large demographic distance from the mainland populations, we cannot distinguish between drift and adaptation. The PCA analysis suggested that drought tolerance and growth may have evolved in a correlated manner. Populations that evolved towards a slower growth were more resistant to water stress, while populations that evolved towards a large stature were the least resistant to water stress (Fig. 4e). Additional evidence for a correlated evolution is that drought tolerance was influenced by a similar set of environmental variables as the growth traits (Fig. 3). Finally, the PCA analysis also suggested that spring phenology evolved independently from the growth-drought tolerance trait complex (Fig. 4e), which is also supported by the fact the two groups of traits seem to have responded to different environmental cues (Fig. 3).

Finally, we were able to corroborate the evidence for adaptive divergence in drought tolerance by comparing observations on seedlings and adults. We estimated drought tolerance in adults using *δ*^13^*C*, a proxy for water use efficiency. We found that the populations’ median *δ*^13^*C* was strongly correlated with seedling’s mean additive trait value for the Water Stress Score (Pearson’s correlation of −0.638, p-value = 0.0078, Fig. 5). As a result, adult trees of populations with a high water use efficiency (WUE) had seedlings that were more resistant to the water stress treatment, i.e. had lower Water Stress Scores, in common garden greenhouse conditions. The main environmental drivers of drought tolerance in seedlings were soil water capacity (PC3) and, to a lesser extent, mean temperature (PC2) (Fig. 3, Table 3). Fig. 5 also illustrates, Available Water Capacity at 30m, which had the highest loading on the synthetic environmental variable, soil water capacity (PC3) (Table 3). Indeed, even a simple visual inspection of Fig. 5 reveals that populations with low Available Water Capacity tend to have more drought tolerant populations as indicated by adult trees’ high WUE and seedlings’ low Water Stress Score (i.e. “light colors go together”).

**Fig. 5:**
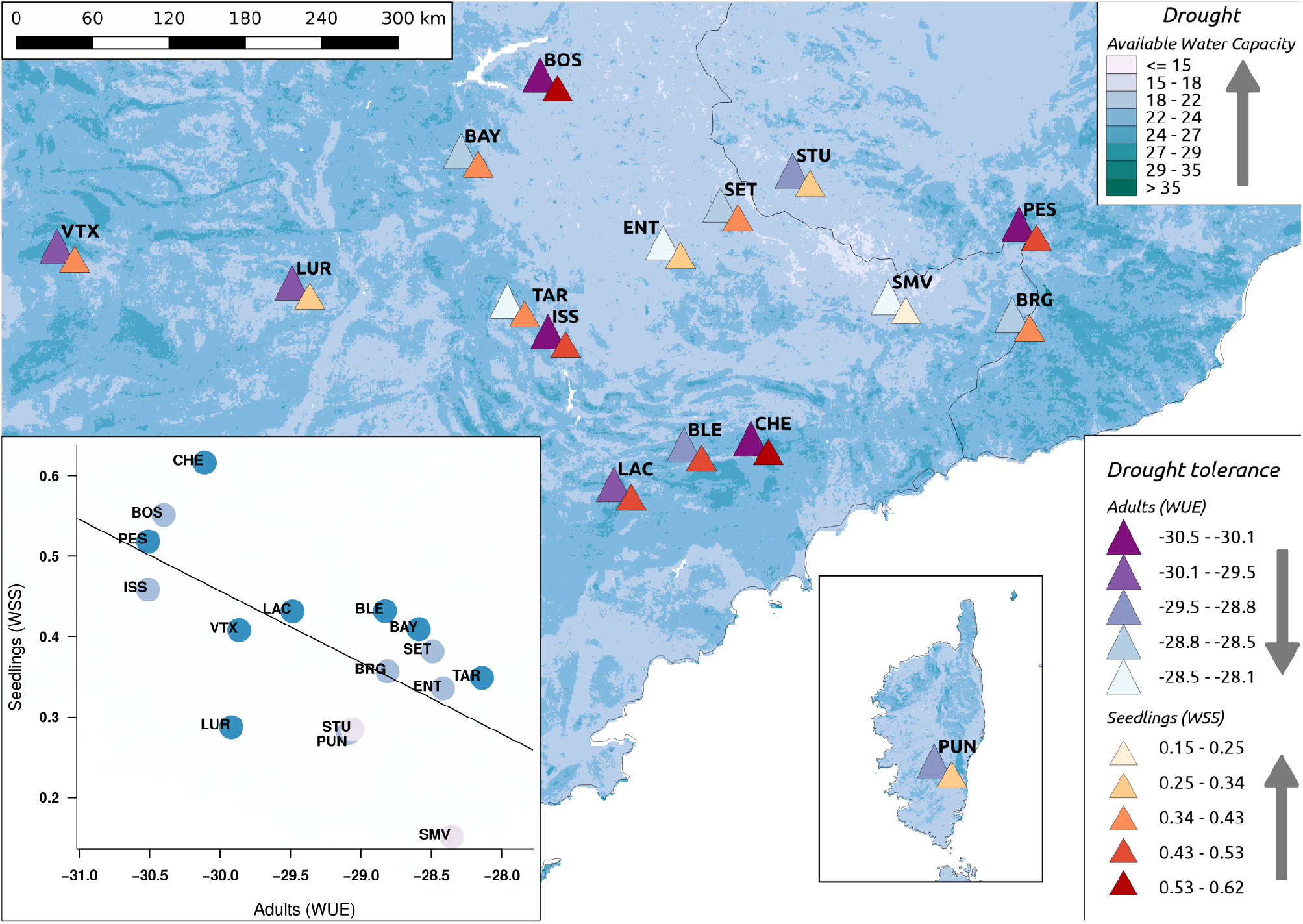
Adaptive landscape of drought tolerance across 16 silver fir (*Abies alba* Mill.) populations based on evidence from seedlings and adults (key entitled “Drought tolerance”). The background of the map shows one of the main environmental drives of adaptation to drought, Available Water Capacity at 30 cm (key entitled “Drought”). Each population is represented by two triangles, showing the estimated drought tolerance of the population based on evidence from adults (large triangles, *δ*^13^*C* referred to Water Use Efficiency (WUE), measured *in-situ*) and seedlings (small triangles, additive genetic trait values of Water Stress Score (WSS), measured in a common garden). The insert scatter plot shows the correlation between adult WUE and seedling WSS; points are color coded for Available Water Capacity at 30 cm.

## Discussion

### Drought tolerance at the southern range margin

We found evidence for adaptive divergence between silver fir populations from the southern margin of the species distribution range for drought tolerance and growth traits. Further, we identified a trade-off between growth and drought tolerance: seedlings originating from populations with relatively slow growth and the small stature were the most drought tolerant both in terms of adult and seedling traits (Fig. 4, Fig. 5). Plant ecophysiologists have long postulated that there is a trade-off between the capacity of genotypes to grow when resources are abundant and to tolerate resource shortages (Bazzaz & Bazzaz, 1996). In arid environments, this paradigm predicts a trade-off between growth potential and the capacity to tolerate drought. This hypothesis has not always been supported by experimental data focusing on interspecies comparisons (*e.g*. Fernández & Reynolds, 2000), including trees (*e.g*. Sack, 2004), potentially due to niche differentiation. In contrast, a growth and/or social status (i.e. dominant or not) versus drought tolerance trade-off has widely been reported at both between- and within-population levels for adult trees (*e.g*. Bennett *et al*., 2015; Grote *et al*., 2016). The underlying physiological mechanism is most likely that water transport to the canopy involve higher cost with increasing tree size, and tall trees experience increased risk of xylem embolism because tension in the water column increases with height (Koch *et al*., 2004; Ryan *et al*., 2006). A growth-drought tolerance trade-off can also be detected in inter-annual growth differences from tree rings: slower growth during drought periods is often observed (*e.g*. Camarero *et al*., 2013; Vitali *et al*., 2017). Finally, some seedling studies also found a growth-drought tolerance trade-off. For example, in Scots pine, Cregg & Zhang (2001) found that seedlings from the drier seed sources survived longer under drought and had a higher *δ*^13^*C*, but were also smaller and allocated more biomass to roots. Moreover, a correlation between seedling growth and adult *δ*^13^*C* was observed across other silver fir populations. Csilléry *et al*. (2018) found that populations from dry inner-Alpine valleys in Switzerland had a higher water use efficiency, and also their progeny grew more slowly in a common garden.

The main environmental driver of drought tolerance and growth were soil water capacity and mean temperature (Fig. 3 and Fig. 5). The result that a soil property is one of the main drivers of selection is comforting. With the Ovaskainen *et al*. (2011) method we inferred the signature of selection on trait optimums in the past, and since soil is more stable than climate, it seems a plausible selective agent. Indeed, forest trees, including silver fir, have large effective population sizes and long generation times, so evolution is slow in comparison to climate fluctuations. A recent study also suggested that soil is more important than climate in determining the distribution of several trees species across Switzerland, including silver fir (Walthert & Meier, 2017).

### Phenotypic plasticity in spring bud phenology

Spring phenology is generally determined by a combination of three key climatic factors, chilling, photoperiod and temperature, among which photoperiod has the strongest genetic component (Körner & Basler, 2010). With the latitudinal range of the studied populations, the photoperiod can be considered constant, thus little genetic variation is expected for bud break phenology. Indeed, we found weak evidence for adaptive divergence in bud break approximated using a Bud Break Score (Fig. 4), which suggests the role of phenotypic plasticity. The main environmental driver of trait divergence in bud break was temperature variance at different temporal scales (daily, seasonally and yearly) and drought variables, such as the frequency and severity in potential evapotranspiration indexes (Table 3, Fig. 3). Weak adaptive population divergence driven by climate variance suggests the role of phenotypic plasticity in the evolution of spring phenology. Phenotypic plasticity occurs when individuals of a given genotype adjust their phenotype according to the conditions they experience (West-Eberhard, 2003). Numerous studies in forest trees have reported that trees possess high levels plasticity and are able to respond phenotypically to a change in their local climate (Kramer, 1995; Alberto *et al*., 2013).

Adaptive divergence in bud break was independent of that in the growth-drought tolerance trait complex (Fig. 4e). As we argued above, many aspects of drought tolerance can be limited by tree stature; i.e. by a characteristic that cannot be changed from year to year. In the past decades, an increase has been observed in the growing season, largely due to earlier springs as a result of increasing temperatures, globally (*e.g*. Linderholm, 2006), and in European forests (Menzel *et al*., 2006). However, forest trees may not be able to benefit from this if there is a shortage in the water supply. For example, Eilmann *et al*. (2011) studied mature Scots pine trees in a dry inner-Alpine valley using an irrigation experiment. They found that control trees, suffering from drought, significant shortened their actual growth period to a much shorter period than the phenological growth period. Further, this result also points towards a possible decoupling of growth and phenology.

### Limitations of the study

The greenhouse experiment itself was a potential source of stress for the seedlings. For example, growing plants in containers of 600 cm^3^ limited the development of their root system, and growing plants in close proximity to each other (distance of 30 cm) created competition for light by the end of the experiment. Additionally, the climate was hotter in Milles (Fig. 1), and the temperature of the greenhouse was more stable than the home environment. The homogenizing nature of the experiment removed much of the environmental variance, which often result in inflated heritabilities in comparison to *in-situ* values. For this reason, the absolute values of the quantitative genetic parameter reported on Fig. 2 should be interpreted only within the context for this experiment. As an example, a recent transplantation experiment of silver fir, also from the French Mediterranean Alps, found that much lower proportion of the trait variance was explained by family than this study (Latreille & Pichot, 2017). Heterogeneity was also introduced via the orientation of the greenhouse: greenhouse 1 was exposed to more wind, which can have a major effect on the temperature of the greenhouse. Mesmoudi *et al*. (2010) found that wind speed staring from 10 m/s may reduce the difference between inside and outside temperature to effectively zero, even at high levels of solar radiation. The effect of this is clearly visible in the results, the mean additive trait values across all populations were different between the greenhouses in 1997, and seedlings grew less, on average, in the “colder” greenhouse (Fig. 4). Nevertheless, it did not affect our conclusions, the populations had similar ordering in terms of trait values and similar degrees of divergence.

Silver fir is a predominantly outcrossing species, thus we assumed that all seedlings from the same mother tree are half-sibs. However, the mating system in silver fir is likely more complicated, and composed of a mixture of outcrossing, bi-parental inbreeding and selfing (Fady & Westfall, 1997). The degree of deviation from outcossing is influenced by a number of ecological factors such as population density. Restoux *et al*. (2008) found that the outcrossing rate and also the variation in outcrossing can be the highest in low density populations, and in silver fir may vary from 0.43 to 0.87. Since we do not have precise information about the stand density or the outcrossing rate, we could not correct for this potential bias. However, since the mating system can have a considerable effect on quantitative genetic parameters (Charlesworth & Charlesworth, 1995), we estimated the *h*^2^ and *CV_A_* by assuming that all seedlings are issued from selfing. Fig. S5 shows that the relative differences in the evolutionary potential and population genetic differentiation would stay similar among traits, however, the absolute values would be three to four times lower. Although most seedlings are probably issued from outcrossing, these values set a lower biological limit on the quantitative genetic parameters in this experiment.

### Conservation and forest management implications

There is abundant evidence of the past evolution of plants in response to climate, such as past range shifts during glaciations (*e.g*. Kawecki & Ebert, 2004; Aitken *et al*., 2008; Ruosch *et al*., 2016). Whether adaptive evolution can compensate for the effects of ongoing human-induced climate change is doubtful. Silver fir may already be in danger of extirpation in the Mediterranean. Ruosch *et al*. (2016) showed that in the French Mediterranean Alps, populations are slowly declining since the last glacial maximum. Recently, die-back events have been documented in the region (Cailleret *et al*., 2014), and also elsewhere at the rear edge of the distribution range. For example, in south-western Europe, reduced growth patterns have been reported in silver fir (Gazol *et al*., 2015). The pace of ongoing climate change is much faster than it has been during the time since post-glacial expansion/re-colonization (Stocker *et al*., 2014). Silver fir may be able to survive in the region by evolutionary rescue (Bell, 2017), or will be taken over by other more drought tolerant species. It has been speculated that late successional tree species, such as silver fir, may lag more behind than early successional species in tracking their climatic niche (Corlett & Westcott, 2013), but the opposite has also been suggested (Dyderski *et al*., 2018). Experimental approaches could provide data to improve such models. For example, using long-term irrigation in a dry inner-Alpine valley, Eilmann *et al*. (2009) could show that facing drought Scots pine is taken over by pubescent oak.

Our results suggest that rapid adaptation to drought conditions could be possible suggested by the high heritability and evolutionary potential for water stress response (Fig. 2). However, we also found that the evolution of drought tolerance is linked to growth (Fig. 5e). A negative genetic correlation between height and drought tolerance rises concerns for current forest breeding practices. Foresters have long been selecting trees for faster growth and height based on provenance trials, and this strategy has been postulated to mitigate climate change effects (Leites *et al*., 2012). Can this scheme also select for drought tolerant genotypes, thus mitigate climate change? Could the evolution of the two traits be decoupled to artificially select for fast growing and drought tolerant genotypes? We think that it is unlikely due to the physiological and biophysical limitation discussed above. For example, Lamy *et al*. (2014) found little plasticity for cavitation resistance, the ability to conduct water through the xylem during drought events. Although Domec *et al*. (2008) found in Douglas fir that structural changes of the xylem could satisfy hydraulic requirements for tall stature, such changes imposed increasing constraints on water transport efficiency. Finally, genetic improvement for both growth and drought tolerance can be possible to an extent that ultimately depends on the genetic architecture of the traits, which is, in practice, often limited by the available genomic resources (*e.g*. Resende *et al*., 2012).

Finally, assisted gene flow can provide a practical solution to mitigate climate change, and could become even a necessity for foundation and resource-production species (Aitken & Bemmels, 2016). Our findings on differences between adaptive strategies among populations may be incorporated in defining assisted gene flow strategies within the study area. Our results are relative to the 16 studied populations, and, it is likely that drought tolerance in all 16 populations is high in comparison to populations in the northern, and more mesic part of the distribution range. Nevertheless, (Csilléry *et al*., 2018) found that dry inner-Alpine valleys in Switzerland had a *δ*^13^*C* that is comparable to that of the studied populations. If the aim was to mitigate climate change and plant southern provenances at higher latitudes, it has to be considered that populations adapted to arid habitats can be at a competitive disadvantage in more mesic environments because of hydraulic and physiological constraints (*e.g*. Eilmann *et al*., 2011; Gessler *et al*., 2017). Monitoring recruitment *in-situ* and measuring water use efficiency in additional populations along a soil water capacity gradient could provide further elements for an adaptive forestry in the region.

## Acknowledgments

The project, and in particular the development of the genetic data, was supported by a research grant from the Center for Adaptation to a changing environment (ACE) at the ETH Zürich and by the group of Alex Widmer (ETH Zürich). KC was supported by an ACE fellowship while collecting the data and by Marie Skłodowska-Curie Individual Fellowship (FORGENET 705972) while analyzing the data and writing this manuscript. We are grateful to the Milles forest nursery team for installing, managing and measuring the experimental plant material. We thank Kristian Ullrich (Max Planck Institute), who helped with the SNP selection, and Dirk Krager (WSL), who helped with extracting the climatic data from the data bases. The contribution of several field and lab workers was necessary for this project, in particular, we thank Olivier Charlandie, who helped with the field sampling, Rene Graf and Fabian Deuber (WSL), who performed most of the sample preparation, and Annika Ackermann (Grassland Sciences Isolab), who performed the stable isotope measurements.

**The authors declare no conflicts of interest.**

## Supplementary Material

**Table S1:**
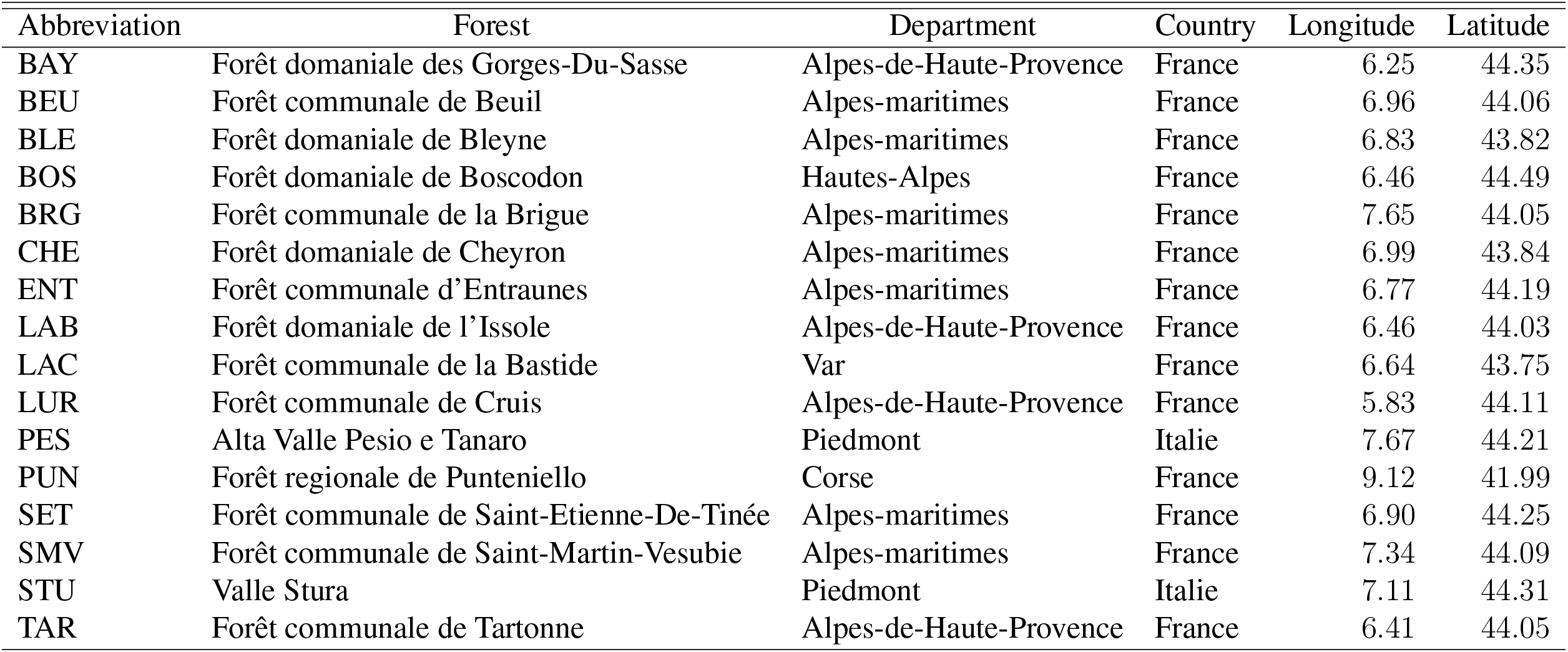
Abbreviated names of the 16 silver fir (*Abies alba* Mill.) population used throughout the paper, the full names of the sampled forests, and their political and geographic situation.

**Table S2:** Geography and environmental variables calculated for the period of 1 January 1901 - 31 December 1978 from monthly mean, minimum and maximum temperature and total precipitation of the CRU TS v. 4.01 data (Harris *et al*., 2014) downscaled using Chelsa data (Karger *et al*., 2017). Soil variables were extracted from the SoilGrids250 data base (Hengl *et al*., 2017). Abbreviations: PET: Potential Evapotranspitation; SPEI: Standardised Precipitation-Evapotranspiration Index.

| Variable | Description | Mean | (Min., Max.) |
| --- | --- | --- | --- |
| **Geography** |  |  |  |
| Long | Longitude (degrees) | 6.9 | (5.2, 9.1) |
| Lat | Latitude (degrees) | 44 | (42, 44.5) |
| Elev | Elevation (m a.s.l) |  |  |
| Slope | Slope (%) |  |  |
| **Standard bioclimatic indexes** |  |  |  |
| bio.1 | Annual Mean Temperature | 6.2 | (3.8, 8.2) |
| bio.2 | Mean Diurnal Range (Mean of monthly Tmax - Tmin)) | 8.4 | (5, 9.5) |
| bio.3 | Isothermality (bio.2/bio.7) (* 100) | 25.1 | (19.6, 26.9) |
| bio.4 | Temperature Seasonality (standard deviation *100) | 611.2 | (508, 649.3) |
| bio.5 | Max Temperature of Warmest Month | 23.1 | (19.6, 25.2) |
| bio.6 | Min Temperature of Coldest Month | -10.3 | (-12.5, -5.8) |
| bio.7 | Temperature Annual Range (bio.5-bio.6) | 33.5 | (25.3, 35.8) |
| bio.8 | Mean Temperature of Wettest Quarter | 2.1 | (0, 6) |
| bio.9 | Mean Temperature of Driest Quarter | 1.8 | (-2.8, 14.4) |
| bio.10 | Mean Temperature of Warmest Quarter | 16.3 | (13.7, 18.3) |
| bio.11 | Mean Temperature of Coldest Quarter | -4.2 | (-6.4, -2) |
| bio.12 | Annual Precipitation | 1163.8 | (801.3, 1671.9) |
| bio.13 | Precipitation of Wettest Month | 396.5 | (271.6, 670.9) |
| bio.14 | Precipitation of Driest Month | 0.3 | (0, 1.6) |
| bio.15 | Precipitation Seasonality (Coefficient of Variation) | 66.6 | (60.6, 77.8) |
| bio.16 | Precipitation of Wettest Quarter | 832.3 | (493.9, 1391.2) |
| bio.17 | Precipitation of Driest Quarter | 35.7 | (11.2, 47.4) |
| bio.18 | Precipitation of Warmest Quarter | 147.1 | (103.8, 204.5) |
| bio.19 | Precipitation of Coldest Quarter | 367 | (107.6, 617.9) |
| **Drought** |  |  |  |
| PET.thorn | Mean annual PET (Thornthwaite) | 43.2 | (37.4, 48.1) |
| PET.harg | Mean annual PET (Hargreaves) | 52.1 | (34, 59.4) |
| SPEI.m1 | Number of month with SPEI < -1 | 147.9 | (142, 153) |
| SPEI.m2 | Number of month with SPEI < -2 | 17.4 | (15, 19.2) |
| SPEI.q5 | 5% quantile of SPEI | -1.6 | (-1.7, -1.6) |
| SPEI.q1 | 1% quantile of SPEI | -2.2 | (-2.4, -2.1) |
| **Late frost** |  |  |  |
| late.frost1 | Min temperature of the first month of the year with mean temperature > 5°C | 1.8 | (1.2, 3.2) |
| late.frost2 | Min temperature of May | 4.2 | (2, 6.7) |
| **Soil** |  |  |  |
| awc10 | AWC (5-15cm) | 49.7 | (44, 53) |
| awc45 | AWC (30-60cm) | 43.6 | (35, 49) |

**Table S3:** Principal Component Analysis (PCA) of 33 environmental variables listed in Table S2. PC axes 1 to 4 explained 88.9% of the variance in the raw environmental variables. Column names show the synthetic names for the PC axes used in the paper, and the variance explained by each. The first ten environmental variables with the highest loadings are shown for each PC axes.

| Temperature variance<br>PC1 (40.1%) |  | Mean temperature<br>PC2 (22.2%) |  | Soil water capacity<br>PC3 (17.8%) |  | Drought<br>PC4 (8.8%) |  |
| --- | --- | --- | --- | --- | --- | --- | --- |
| Variables | Loadings | Variables | Loadings | Variables | Loadings | Variables | Loadings |
| bio.7 | -0.28 | late.frost2 | 0.31 | awc45 | 0.36 | SPEI.m2 | 0.49 |
| bio.2 | -0.28 | PET.thorn | 0.3 | awc10 | 0.29 | SPEI.q5 | -0.35 |
| bio.4 | -0.28 | bio.1 | 0.29 | bio.1 | 0.27 | bio.19 | 0.33 |
| bio.3 | -0.27 | bio.8 | 0.28 | PET.thorn | 0.26 | bio.8 | -0.3 |
| late.frost | 0.27 | bio.10 | 0.27 | bio.17 | 0.26 | bio.9 | 0.28 |
| PET.harg | -0.26 | bio.18 | -0.27 | bio.13 | 0.26 | awc10 | -0.27 |
| SPEI.q1 | -0.26 | bio.11 | 0.25 | bio.12 | 0.26 | bio.18 | -0.26 |
| Y | -0.26 | bio.19 | -0.24 | bio.10 | 0.25 | bio.14 | -0.23 |
| bio.15 | 0.24 | bio.12 | -0.24 | bio.16 | 0.24 | SPEI.m1 | 0.2 |
| X | 0.23 | bio.16 | -0.21 | late.frost2 | 0.24 | Y | -0.16 |

## Supplementary figures

**Fig. S1:**
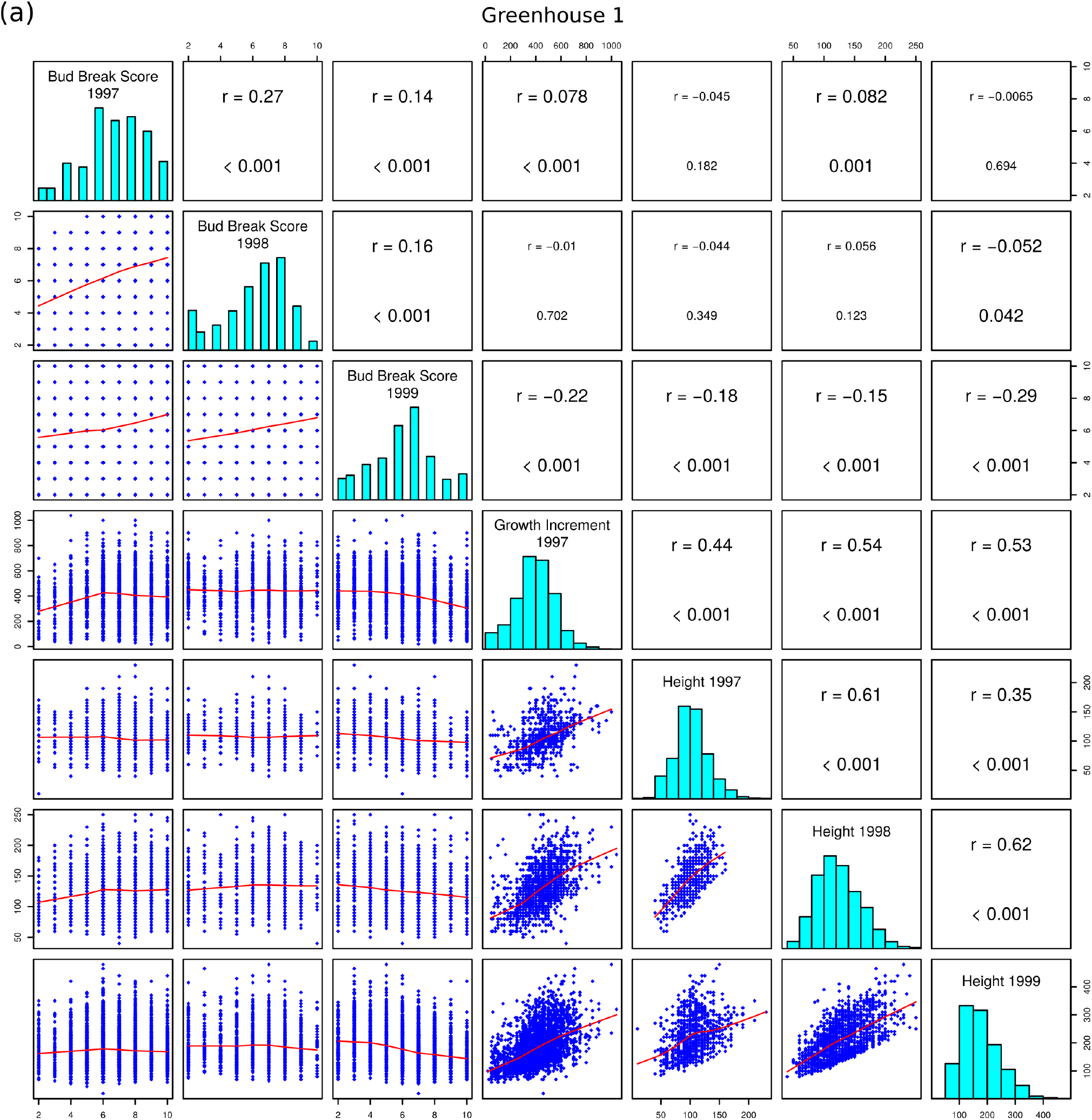
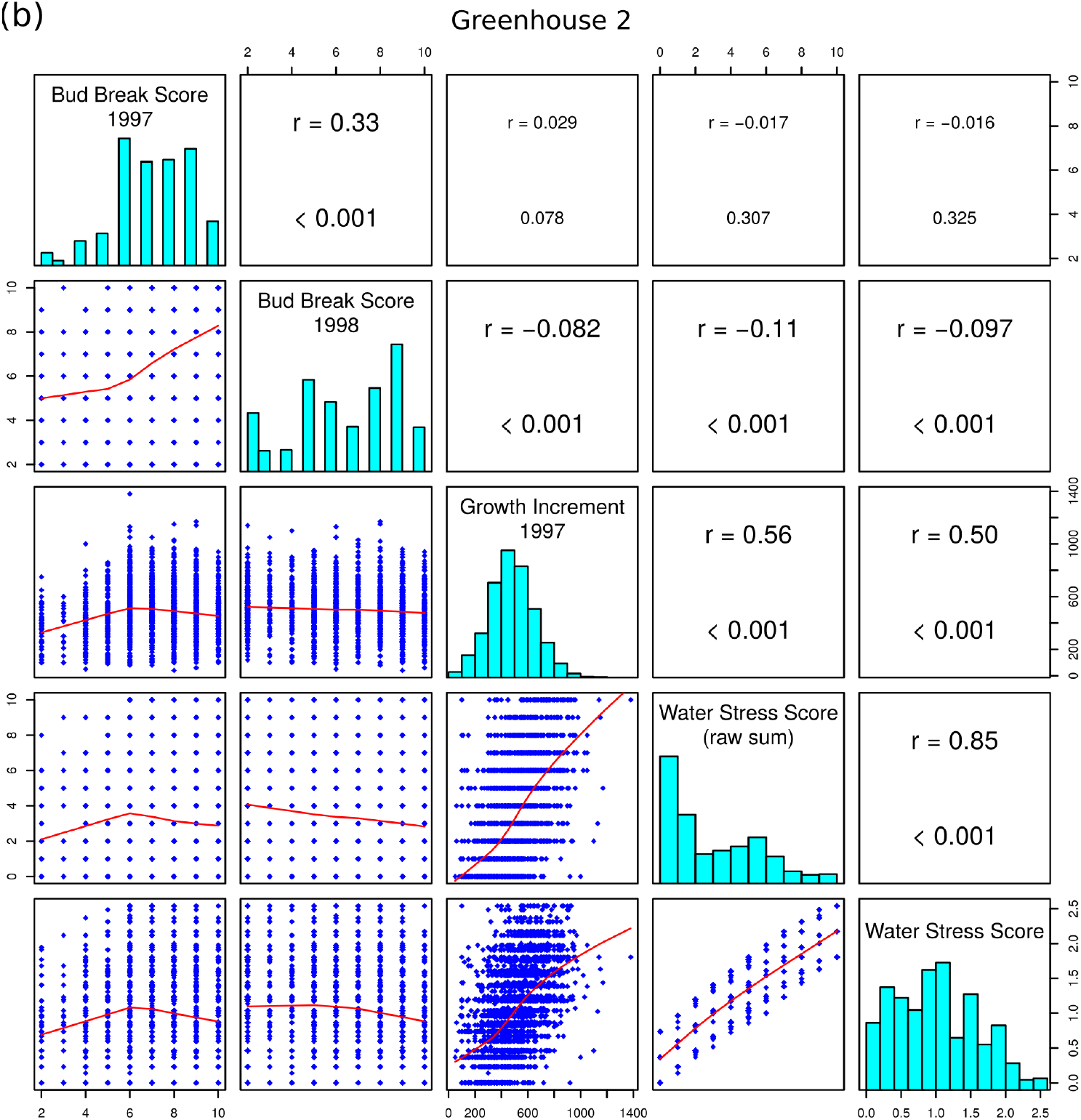
Raw seedling trait values and the correlation between them recorded in greenhouse 1 (a) and greenhouse 2 **(b)**. Seedlings originated from 16 silver fir (*Abies alba*) populations across the French Mediterranean Alps. Panels in the upper triangles show the Pearson correlation and the p-value from the correlation test. Significant correlations (p-value < 0.05) are shown in larger fonts. Panels in the lower triangle show a scatter plot between the two variables with a smooth curve in red fitted using the *lowess* function in R. Panels in the diagonal show the distribution the variables as histograms.

**Fig. S2:**
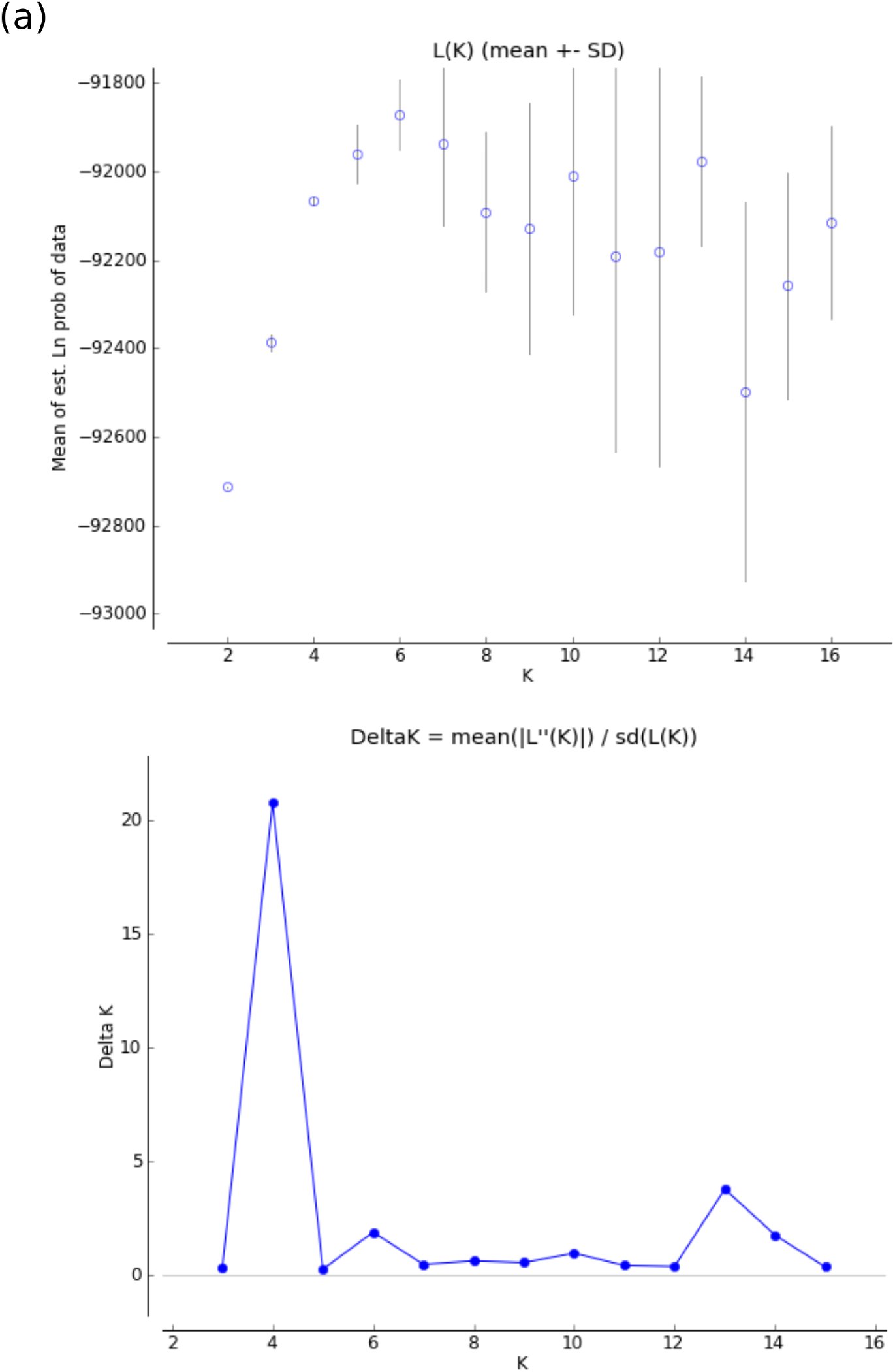
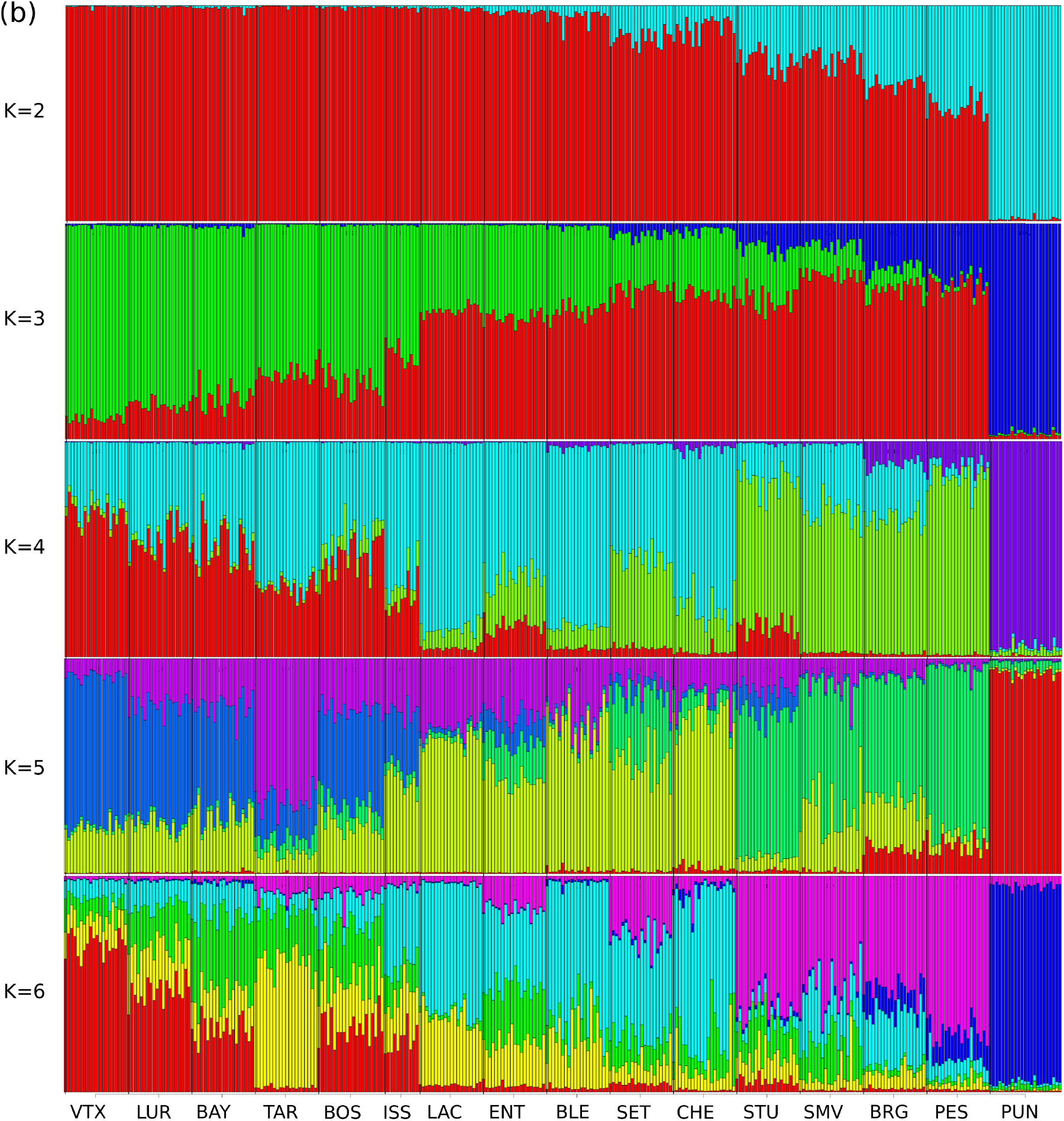
Population genetic structure of 16 silver fir (*Abies alba*) populations across the French Mediterranean Alps inferred using the software *Structure* (Falush *et al*., 2003) using 357 SNP loci. **(a)** Log likelihood of the data and the delta K values (Evanno *et al*., 2005) for K values between two and 16. **(b)** Proportion of ancestry of each sampled tree (colored vertical bars) from each of the assumed genetic clusters (K). Each cluster is indicated with a different color and colors are arbitrary between different values of K. Results for K values between one and six are shown.

**Fig. S3:**
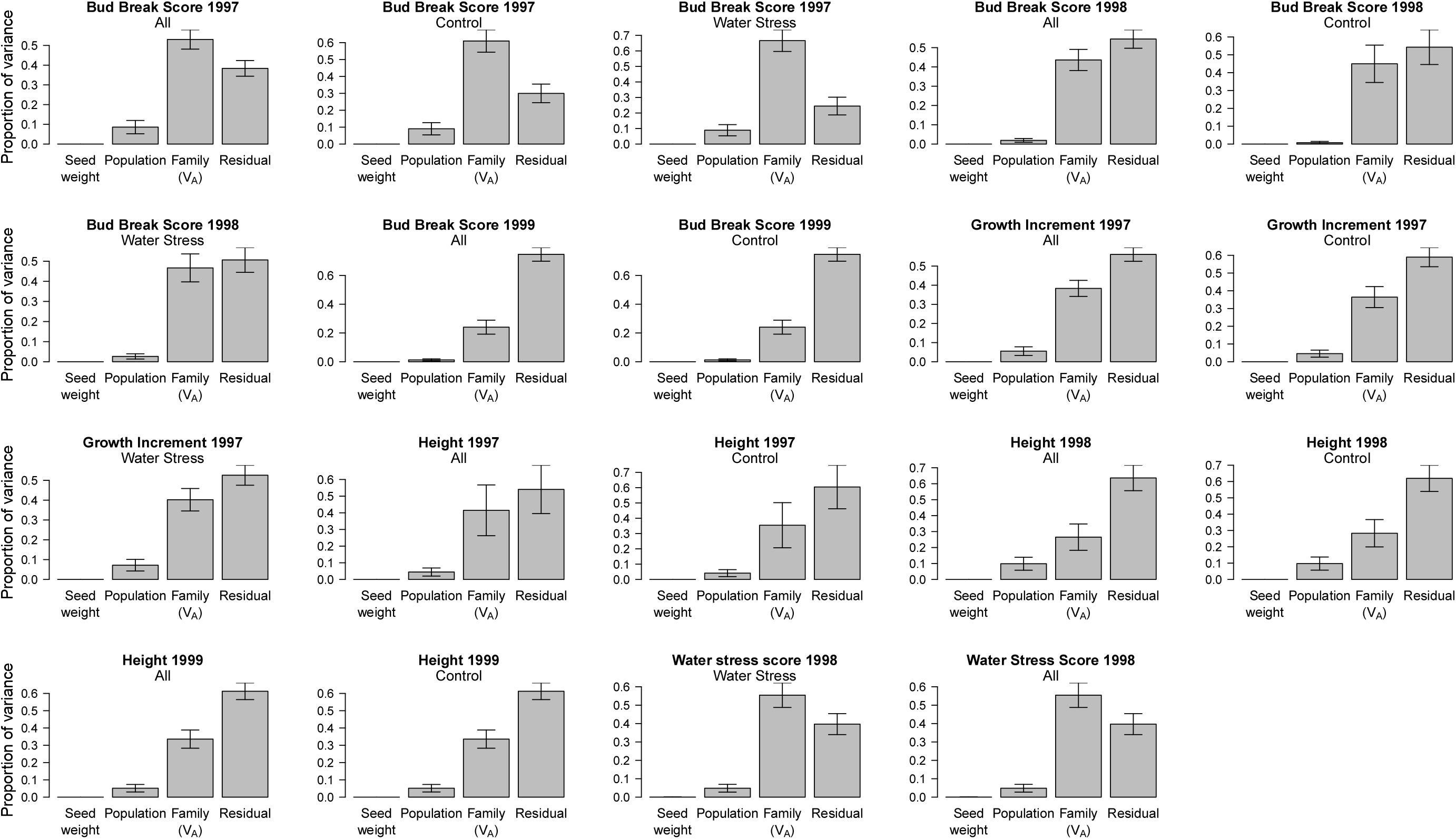
Variance components estimated using a mixed-effects model called the animal model implemented in *ASreml-R* including block as a fixed effect, and seed weight and population, and family structure (i.e. the pedigree) as random effects. *V_A_* stands for the additive genetic variance, which is the trait variance due to resemblance between family members.

**Figure S4.**
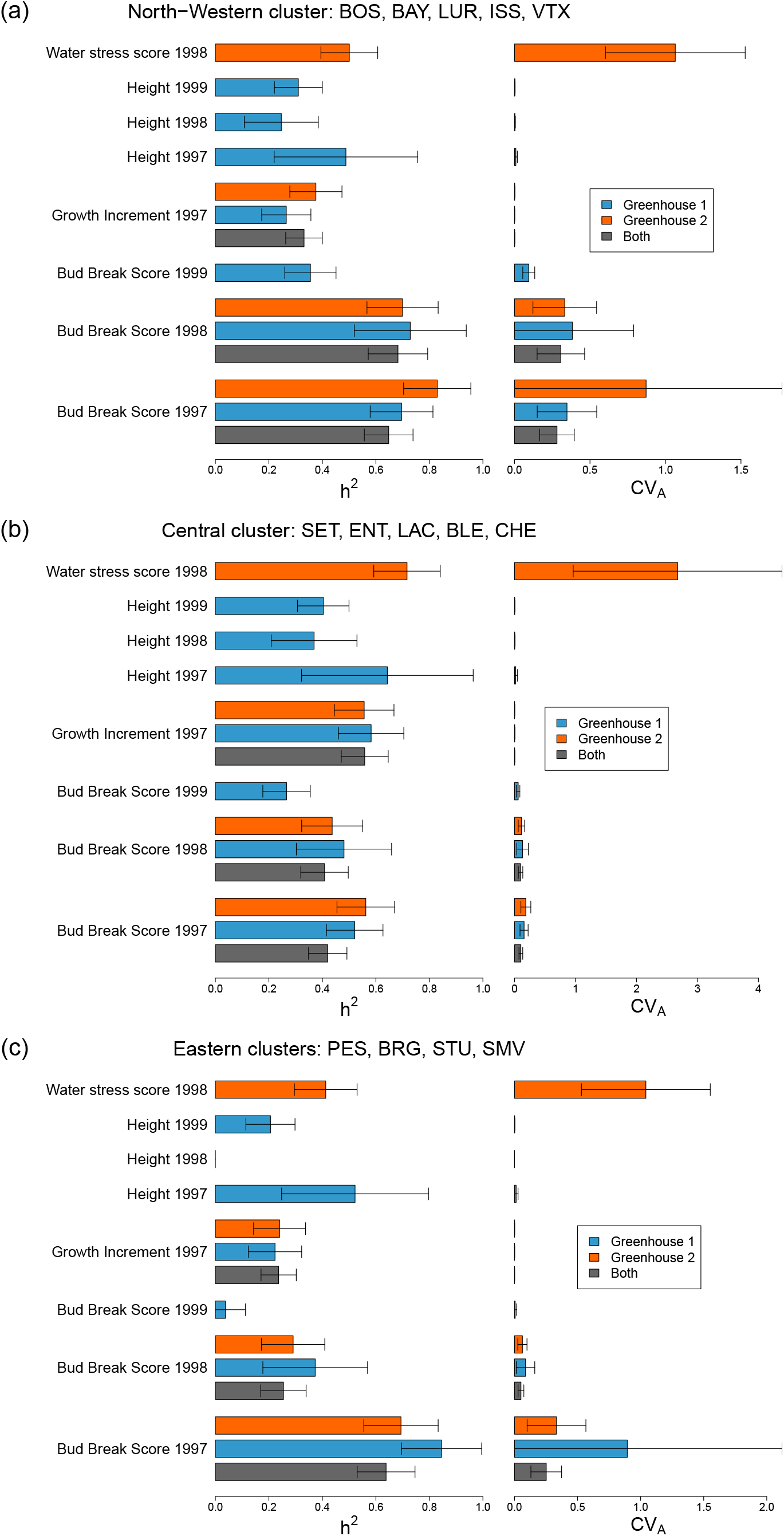
Heritability (*h*^2^) and additive genetic coefficient of variation (*CV_A_*) of 11 traits measured on silver fir (*Abies alba*) seedling in a common garden. The full data set comprising two greenhouses and 8199 observations were divided into three groups according to the genetic and geographic clustering of the populations. **(a)** North-Western cluster (N=2496), **(b)** Central cluster (N=2639), and **(c)** Eastern cluster (N=1883). Two genetically isolated populations, TAR and PUN, were excluded from this analysis. Not all traits were scored in both greenhouses. Greenhouse 2 received a water stress treatment in 1998 after which only Water Stress Score was recorded. Parameters were estimated using an animal model implemented in *ASreml-R* including block as a fixed effect, and seed weight and population as random effects.

**Fig. S5:**
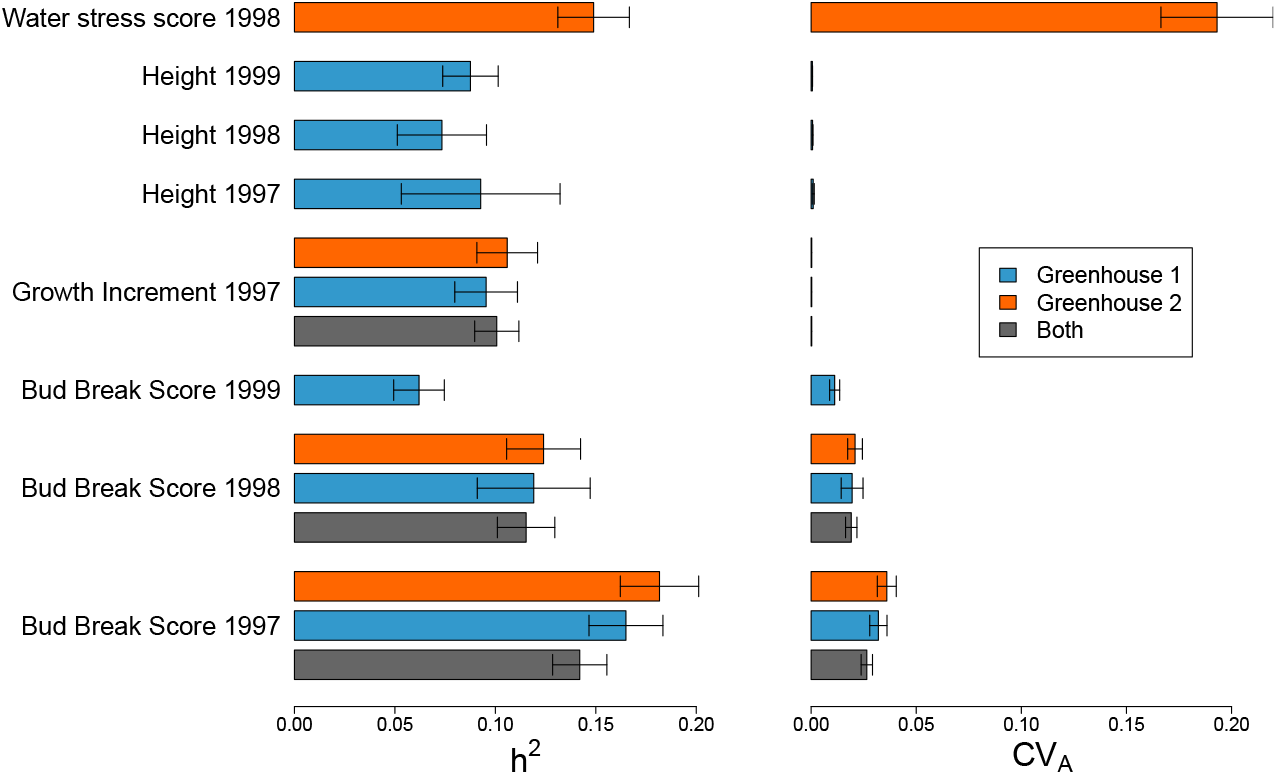
The lower bound on heritability (*h*^2^) and additive genetic coefficient of variation (*CV_A_*) estimated assuming that all seedlings were issued from selfing. 11 traits measured on silver fir (*Abies alba*) seedling in a common garden. The experiment consisted of two greenhouses and counted 8199 observations. Not all traits were scored in both greenhouses. Greenhouse 2 received a water stress treatment in 1998 after which only Water Stress Score was recorded. Parameters were estimated using an animal model implemented in *ASreml-R* assuming including block as a fixed effect, and seed weight and population as random effects.

## Footnotes

1 https://www.helsinki.fi/en/researchgroups/metapopulation-research-centre/rafm-and-driftsel

2 https://www.helsinki.fi/en/researchgroups/metapopulation-research-centre/rafm-and-driftsel

